# helixCAM: A Platform for Programmable Cellular Assembly in Bacteria and Human Cells

**DOI:** 10.1101/2022.04.19.488034

**Authors:** George Chao, Timothy M. Wannier, Clair Travis, Nathaniel C. Borders, Evan Appleton, Anjali Chadha, Tina Lebar, George M. Church

## Abstract

Interactions between cells are indispensable for creating structure and signaling. The ability to direct precise cell-cell interactions would be powerful for engineering tissues, understanding signaling pathways, and directing immune cell targeting. In humans, intercellular interactions are mediated by cell adhesion molecules (CAMs). However, CAMs are natively expressed by many cells and have cross-reactivity, making them unsuitable for programming specific interactions. Here, we showcase “helixCAM,” a platform for engineering novel CAMs by presenting coiled-coil peptides on the cell surface. helixCAMs were able to create specific cell-cell interactions and direct patterned aggregate formation in bacteria and human cells. We built rationally designed helixCAM libraries and discovered novel high-performance helixCAM pairs. High-affinity helixCAMs were then used for multicellular engineering applications, such as spherical layering, adherent cell targeting, and surface patterning. The helixCAMs are an expandable platform for directing complex multicellular assemblies, and we foresee its utility in tissue engineering, immunology, and developmental biology.

## Introduction

Almost all organisms, whether unicellular or multicellular, engage in intercellular interactions. In bacteria, pili facilitate conjugation and the exchange of genetic material[1]. For multicellular organisms, cell-cell interactions dictate a range of important physiological functions, including stem cell differentiation[2], morphological development and maintenance[3], and the activation of immune responses[4].

The ability to program specific cell-cell interactions has numerous applications. For instance, the aggregation of bacterial cells can be used to create living biomaterials with the ability to release antibiotics [5] or undergo self-repair [6]. Programmed cell-cell interactions can be used to study immune response activation in T-cells [7]. Additionally, by using programmed cell-cell interactions to pattern and aggregate cell assemblages, there is the potential to build synthetic tissues [8]. In terms of constructing synthetic tissues using multiple cell types, current approaches primarily rely on nozzle-based cell printing of terminally-differentiated cells [9]– [12]. This method of creating cell structures faces significant obstacles from low cell viability due to shear stress and difficulty for differentiated cells to form junctions after deposition[13], [14]. Rather than the top-down approach of printing cells, allowing cells to form patterned structures through selective binding could resolve both limitations and enable the synthetic construction of more complex, accurate, and viable tissues.

Some approaches exist to artificially direct cell-cell interactions. By overexpressing two cadherins in CHO cells, Shan *et al.* were able to assemble large cell aggregates[15]. Native cell adhesion molecules (CAMs) such as cadherins, however, are finite, have significant cross-activity [16], and often can play dual roles as downstream signaling molecules[17], limiting their use for directing multiple interactions in parallel. Another approach is to conjugate the cell membrane with single-stranded DNA (ssDNA), allowing cells to selectively interact via base-pairing [18]. In this case, limitations arise in the stability of extracellular DNA[19] as well as the need to chemically treat each cell population, preventing the control of cell interactions through genetic programs or small molecule inducers. A genetically-encoded mechanism to control cell-cell interactions was recently demonstrated in *E. coli*, using surface-mounted nanobodies and antigens [20]. This groundbreaking approach enables the collection of known nanobody-antigen pairs to be repurposed for programming cell-cell interactions but still comes with some drawbacks from limited expandability and the potential for unintended biological responses to the surface-presented nanobody or antigen.

One attractive alternative for protein-mediated cell interaction is coiled-coils (CCs), well-studied protein domains with established roles in mediating protein-protein interactions across numerous native proteins[21]. Each CC consists of multiple seven amino-acid repeats, termed “heptads,” which chain together to form alpha-helical secondary structures[22]. The specific residues forming the heptads determine the orientation, binding multiplicity, and specificity of each CC [23]. The potential of these short peptides to mediate interactions between micron-scale objects has been demonstrated by Obana *et al.*, who aggregated polystyrene beads functionalized with CCs[24]. Moreover, due to their well-understood pairing rules, collections of orthogonal CC peptide pairs have been engineered and characterized by numerous research groups[25]–[28], making them a potential treasure trove for inducing specific cell-cell interactions.

We hypothesized that presenting CC domains fused to a transmembrane domain (TMD) on the surface of cells may create tight and programmable cell-cell interactions. We term this general CC-TMD design paradigm “helixCAM” (Figure 1A). Within this work, we find that helixCAMs direct selective binding of bacteria and human cells into patterned aggregates and engineer novel CC peptides for helixCAM use, for a total of five pairs. Using these, we implement helixCAMs for a wide range of applications, such as specifying the spatial composition of spherical aggregates, targeting suspension cells to adherent cells, and patterning helixCAM cells with His-tagged CC peptides. The helixCAM platform is a foundational technology that empowers researchers with simultaneous control of multiple specific cell interactions. We expect scientists and engineers across biological disciplines to find compelling applications for helixCAMs in their work, and for the set of orthogonal helixCAMs to grow and open the door to programming increasingly complex interactions.

**Figure 1.**
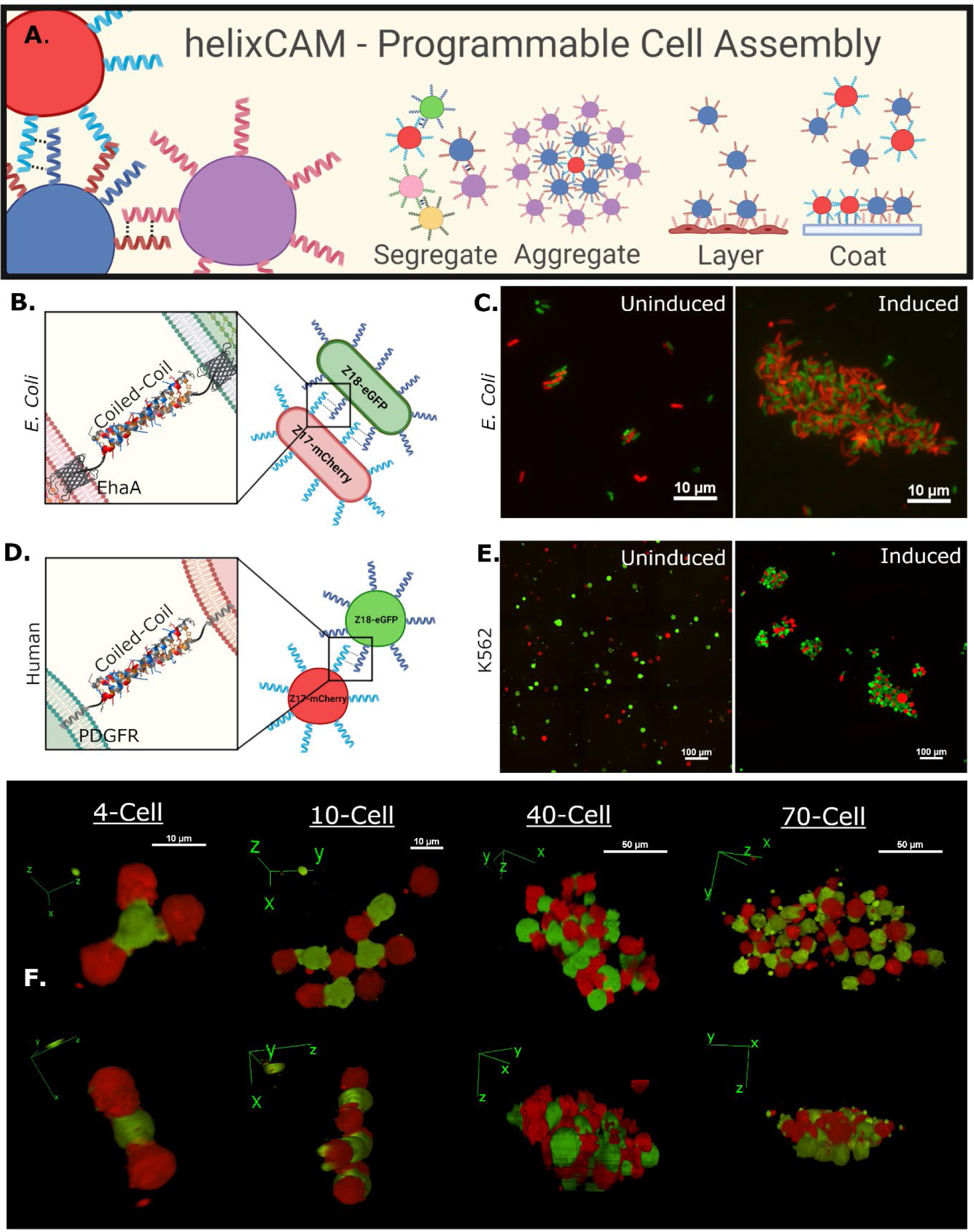
Overview of helixCAM design and applications and demonstration with Z17/Z18. **A.** Schematic of the helixCAM concept and potential applications. **B.** Design of helixCAMs in *E. coli*: the coiled-coil domain is fused to the EhaA autotransporter protein for surface presentation (Details in S1). **C.** helixCAM-induced *E. coli* aggregation, with Z17 cells co-expressing mCherry, and Z18 cells co-expressing eGFP. Representative views are shown from several fields of view. Images were taken at 60X magnification and cropped (uncropped images in S2). **D.** Design of mammalian helixCAMs: the coiled-coil domain is fused to the PDGFR transmembrane domain (Details in S1). **E.** helixCAM-induced K562 aggregation, with Z17 cells co-expressing mCherry, and Z18 cells co-expressing eGFP. Representative views are shown from several fields of view. Image was taken at 20X magnification with 3x3 tiling and cropped (uncropped images in S2). **F.** 3D reconstruction of helixCAM-induced K562 aggregates of various sizes, demonstrating heterodimeric three-dimensional patterning. Images were taken at 40X magnification using a spinning disc confocal system as a Z-stack and reconstructed using ImageJ.

## Results

### helixCAMs induce large, predictably-patterned multicellular aggregates in *E. coli*

To pilot helixCAMs as a cell-cell interaction platform, the antiparallel heterodimeric CC pair Z17 and Z18 was selected from the SynZip library [25]. To generate helixCAMs in *E. coli*, CCs were fused to the N-terminus of the *E. coli* EhaA autotransporter adhesin protein [29] (Figure 1B, S1), and expressed under the IPTG-inducible T7lac promoter[30]. We tested these bacterial helixCAMs in two distinct strains: one expressing the Z17 helixCAM along with the mCherry fluorescent protein, and another expressing the Z18 helixCAM alongside GFP. When the two populations were mixed without arabinose induction, bacterial cells were mostly singlets, whereas with arabinose induction we observed formation of large bacterial aggregates. The images shown in Figure 1C are representative of aggregates observed across multiple fields of view (uncropped image can be found in S2). Interestingly, the cells formed a distinctive alternating pattern between mCherry- and GFP-expressing cells, demonstrating the desired heterodimerizing effect.

### HelixCAMs are amenable to human cell line programming

We next sought to adapt the helixCAM system in human cells. To this end, we inserted the Z17 and Z18 CCs at the N-terminus of the platelet-derived growth factor receptor transmembrane domain and C-terminus of the secretion signaling peptide from human Immunoglobulin K, following the design from Chesnut *et al.* (Figure 1D) [31]. Expression of human helixCAMs was placed under the Tet-On doxycycline-inducible promoter[32] along with constitutive expression of an identifying fluorescent protein under the EF1α promoter[33] and flanked with PiggyBac sites for stable line generation [34] (S1).

We then generated stable cell lines for each helixCAM in human K562 cells, an immortalized leukemia cell line[35] selected for their lack of innate cell adhesion and spherical shape conducive to efficient packing. Similar to our experiments in *E. coli*, we tested the capability of helixCAMs to aggregate human cells by mixing the Z17^mCherry^ cells with Z18^eGFP^ cells. Without induction, cells remained as single-cell suspensions, but after 48 hours of induction, cells formed large, tight clusters with an alternating pattern of mCherry and GFP indicative of heterodimerization (Figure 1E). Visualizing the three-dimensional structure from confocal imaging reconstructions (Figure 1F), we observed that helixCAM proteins induced junction-like binding interfaces between cells. Smaller aggregates of four to ten cells formed structures in which each cell bound exclusively to cells expressing the complementary helixCAM, forming clear checkboard patterns. At higher cell counts, the alternating pattern is present but less precise, and aggregates spanned multiple cell layers in X, Y, and Z directions, forming three-dimensional aggregates spanning hundreds of microns in each direction. These data demonstrate that helixCAMs can form strong cell-cell interactions in human cell lines independently of endogenous CAM proteins.

### Orthogonal helixCAM pairs enable programmable sub-aggregation within a mixed population

From the promising results of Z17/Z18, we designed four additional helixCAMs using the CC pairs P3/AP4 and P9/AP10, which were previously reported to have high affinity and orthogonality [26], [28]. We first tested these new helixCAMs in *E. coli* using a sedimentation rate assay[36] and observed increased settling when mixing *E. coli* expressing the intended CC pairs, indicating the formation of larger aggregates and thus higher CC affinity (S3). To test these new helixCAMs in human cells, we generated four additional helixCAM K562 lines and denoted each helixCAM line with a corresponding fluorescent protein (P3^iRFP670^, AP4^eBFP2^, P9^mPlum^, and AP10^mOrange^). As in the case of Z17/Z18, both P3^iRFP670^/AP4^eBFP2^ cell lines (Figure 2A) and P9^mPlum^/AP10^mOrange^ cell lines (Figure 2B) form cell aggregates when induced to express helixCAMs in paired co-cultures.

**Figure 2.**
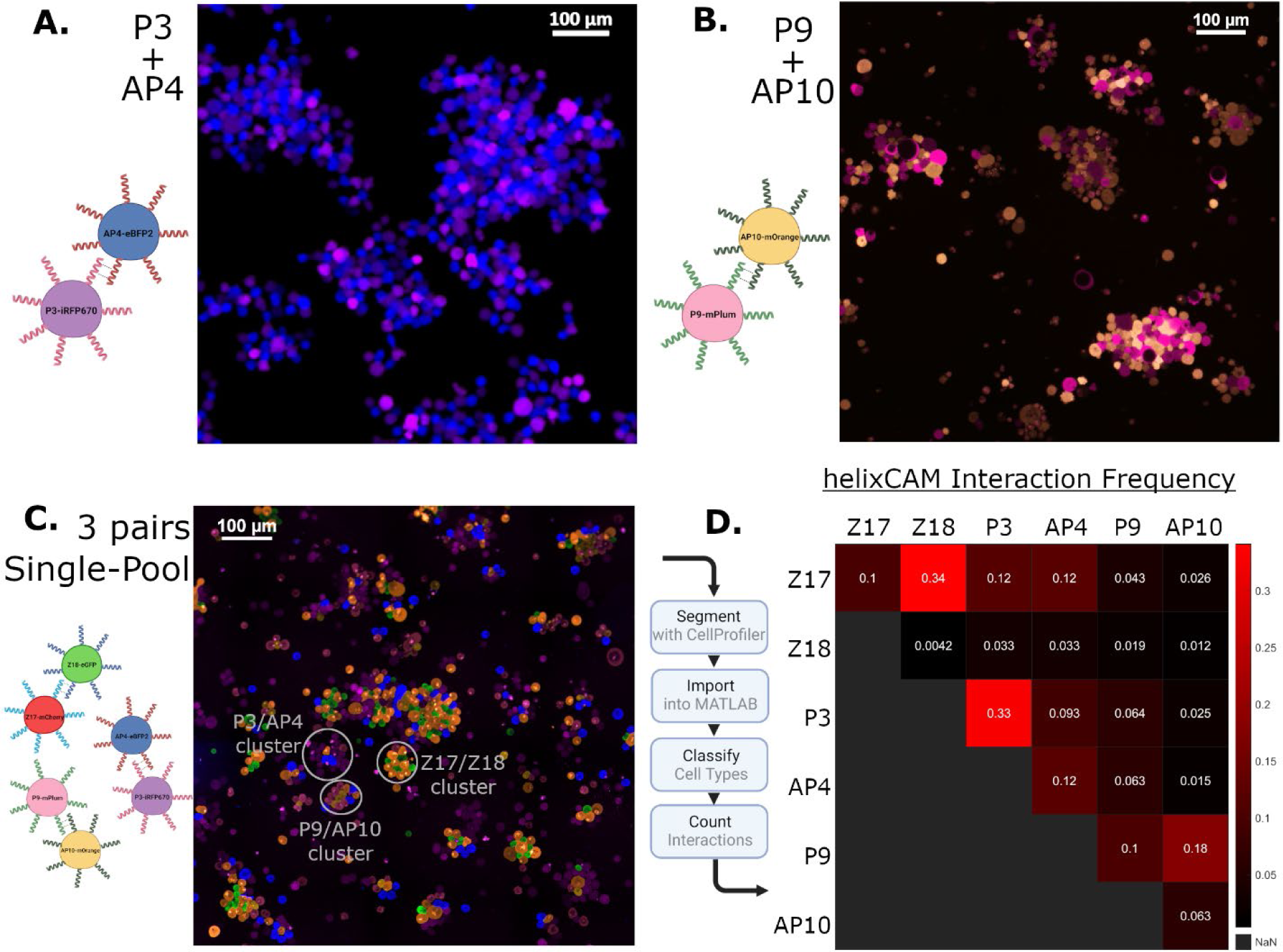
Characterization of additional helixCAM pairs and three-pair interaction orthogonality. **A.** K562 cell aggregates formed by P3/AP4 interactions. P3 cells co-express the iRFP670 and Z18 cells co-express eBFP2. Image was taken at 20X magnification and cropped (uncropped image in S2). **B.** K562 cell aggregates formed by P9/AP10 interaction. P9 cells co- express mPlum, and AP10 cells co-express mOrange. Image was taken at 20X magnification and cropped (uncropped image in S2). **C.** Z17^mCherry^, Z18^eGFP^, P3^iRFP670^, AP4^eBFP2^, P9^mPlum^, and AP10^mOrange^ cells were induced to express helixCAMs in a single mixed culture. Clear sub- clusters can be observed. Image was taken at 20X with a 6x6 tile across five channels and cropped to show regions of interest (uncropped image in S4). **D.** Interaction frequency table was derived from the uncropped three-pair co-culture image. (Detailed in S5).

Next, we sought to examine the orthogonality of these interactions when all three pairs are present in a single culture. To this end, the six cell lines were seeded together at identical concentrations and helixCAM expression was induced for 48 hours. Microscopic images of the resulting multicellular assemblies showed clear binding preferences between complementary helixCAM cells (Figure 2D). Individual aggregates comprised primarily of pairs of complementary helixCAM cells, and, while larger aggregates were composed of more cell types, indicating some promiscuous binding, distinct sub-clusters were still evident.

To quantify the orthogonality of the interactions in the structures, we performed image segmentation of a six-by-six field of view (S4) and estimated the frequency of interaction between each of the six cell types (Figure 2D). Four cell lines, Z17^mCherry^, Z18^eGFP^, P9^mOrange^, and AP10^mPlum^, showed a clear preference for binding to their complementary partners. However, while the Z17^mCherry^ cells bound complementary Z18^eGFP^ cells at the highest frequency, off-target interactions were also observed with P3^iRFP670^ and AP4^eBFP2^. The P9^mOrange^ echoed this trend to a lesser extent, with the highest frequency of interactions with complementary AP10^mPlum^ cells along with a handful of off-target interactions with P3^iRFP670^ and AP4^eBFP2^. Notably, the behavior of helixCAMs in human cells contrasted with the higher orthogonality observed in *E. coli*. (S3). This gap suggested that protein expression and transport may be factors in the ultimate helixCAM function and indicated a need for a eukaryotic screen when engineering CCs for use in helixCAM applications.

### Rational design and two-stage screening of helixCAM library yields two novel, high-performance helixCAM pairs

To build more complex cellular structures, we sought to expand the set of high-affinity and orthogonal helixCAMs by designing rationally-designed CC peptide libraries based on the four helixCAMs that exhibited high affinity and specificity in K562 cells (Z17, Z18, P9, AP10). As mentioned above, CCs are composed of heptads, within the seven amino acid positions designated as “a” through “g” [21]. As illustrated in Figure 3A, electrostatic interactions in positions “g” and “e” determine binding specificity, whereas hydrophobic residues in positions “a” and “d” interact at the binding interface to affinity and stability [37], [38]. To introduce variability in heptad interactions, we placed either glutamic acid or lysine residues in the “g” and “e” positions, with the rationale that opposing charges in these positions will increase the likelihood of generating novel pairs with little specificity to the template CCs. Additionally, we placed either hydrophobic (leucine/isoleucine) or polar (asparagine) residues in the core binding region (“a” and “d”) to seed variability in baseline affinity. Each CC utilized for helixCAM consists of five heptad repeats, and we combinatorially mutate three residues per repeat to one of two amino acids, leading to a library size of 2^15^, or 32,768 members, for each template. Libraries were screened against themselves and all other libraries, resulting in 10 paired libraries, and approximately 10 billion possible interactions. To effectively screen these interactions, we employed a two-stage approach: first, a tripartite split-GFP complementation assay was performed in *E. coli* to select for CCs candidates exhibiting strong binding. This primary screen was followed by a yeast mating assay to evaluate CC pairs for their ability to induce eukaryotic cell aggregation.

**Figure 3.**
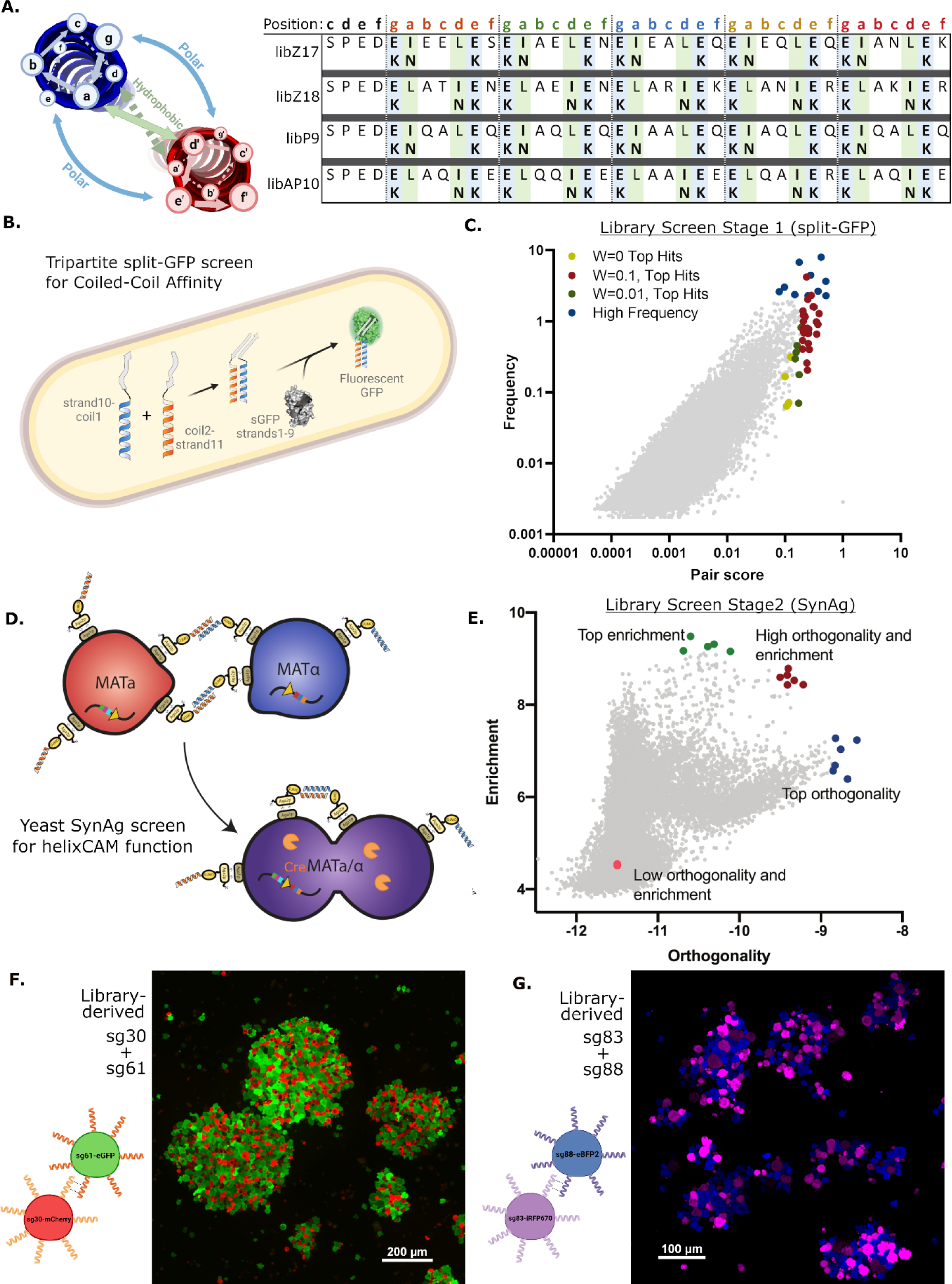
Rational Design of helixCAM-optimized Coiled-Coil library and two-stage screen for helixCAM performance. **A.** Table of amino acid substitutions for each of four template-derived helixCAM libraries. Each template consists of five heptads, within which either the “g”, “a” and “e” position or the “g”, “d”, and “e” positions are varied to form new electrostatic and hydrophobic interactions. **B.** Design of the tripartite split-GFP assay for CC affinity. One CC library was fused to β-strand 10 and another to β-strand 11 of the tripartite split-GFP. Interaction between CCs stabilizes a fluorescent GFP protein. **C.** Graph of CC candidate pairs’ frequency in the population versus their pair score (graph uses W=0.1). Each point represents a pair of CCs. The pair score represents a pair’s specificity to each other and is adjusted with a weight parameter W (S7). The top 30 hits using three different W’s, along with several high-frequency pairs (totaling 102 individual CCs) were selected for subsequent screening. **D.** Design of modified yeast SynAg mating assay for helixCAM-compatible CC selection. Haploid yeast cells MAT**a** or MATα expressing surface-presented CC candidates were mixed. CC binding induces haploid cells to mate, creating a diploid cell that survives dual auxotrophic selection. The fusion of cells also leads to the expression of the Cre recombinase (orange), which integrates the two CC constructs and their barcodes into the same DNA strand. **E.** Stage 2 CC candidate pairs’ enrichment versus their orthogonality. Enrichment is the observed frequency of the pair, whereas orthogonality is the frequency of the pair divided by the total observations of each of the two CCs in the pair. Two pairs, sg30/sg61 and sg83/sg88, were selected from the red group for helixCAM use. **F.** Large aggregates of sg30^mCherry^ and sg61^eGFP^. Image was taken at 20X magnification with a 3x3 tile and subsequently cropped. G. Aggregates of sg83^iRFP670^ and sg88^eBFP2^. Image was taken at 20X magnification and presented without cropping.

The tripartite split-GFP method was selected for its low background and high throughput readout via FACS [39], [40]. In our assay, sGFP β-strands 1-9 were constitutively expressed, β- strand 10 was fused to the C-terminus of one CC library, while β-strand 11 was fused to the N-terminus of a second CC library (Figure 3B). All three components were combined into a single plasmid and transformed into *E. coli* cells as pairwise mixtures. Cells with high GFP signals were sorted (S6), and the selected plasmids were sequenced with next-generation sequencing (NGS) to identify candidate CC pairs. Each pair was assigned a “pair score,” which positively weighed the frequency of on-target binding (as a metric of affinity) and negatively weighed the frequency of off-target binding. Using this score, 102 novel CC peptides were selected for the next stage of the screen (Figure 3C).

Considering the potential disconnect between prokaryotic and eukaryotic helixCAM characteristics, the next stage of CC screening was designed to select for the ability to induce aggregation in eukaryotes while maintaining a reasonable screening throughput; the yeast-based SynAg mating assay was a good fit [41]. Briefly, MAT**a** and MATα haploid yeast cells, respectively carrying lysine or leucine auxotrophic markers, were transformed with the yeast helixCAM library (S8) comprised of the 102 CC candidates. The two mating types are then incubated in a shaking culture. Yeast cells displaying interacting helixCAMs were promoted to mate, merging the two haploid cells into a diploid cell. Diploid cells gain both auxotrophic markers and are positively selected. Merged diploid yeast cells also began to express the Cre recombinase, which recombined the two helixCAM DNA constructs into a single strand (Figure 3D). After using NGS to examine the barcodes of yeasts that survive selection, we assigned an “enrichment” and an “orthogonality” score for each helixCAM pair and, using these metrics, selected two novel CC pairs (sg30/sg61 and sg83/sg88) that were likely to be capable of directing strong and specific cell aggregation (Figure 3E).

To test the new CC pairs, we built four K562 cell lines that inducibly express the library-derived helixCAMs and repeated the pairwise aggregation experiment (Figure 3F, 3G). We found that both new helixCAM pairs robustly generated patterned cell aggregates. In particular, the sg30/sg61 pairing consistently yielded aggregates containing thousands of cells, and some aggregates spanned millimeters in length (S9) and were visible by eye (S11).

### “Human cell sedimentation assay” reveals high affinity and specificity in library-derived helixCAMs

With the addition of the two novel helixCAM pairs, we had ten distinct helixCAMs to test for binding orthogonality. As it is challenging to distinguish beyond eight fluorescent proteins through conventional fluorescent microscopy, an imaging-independent assay was required to determine pairwise affinity across all helixCAMs. Using the same principles of the *E. coli* sedimentation rate assay, we developed a human cell sedimentation rate assay (HCSRA) to assess aggregation in K562 cells. For this assay, helixCAM-presenting cells were co-cultured in ultra-low-adhesion V-bottom plates for 48 hours and evaluated for settling by measuring cell settling over time through optical density (Figure 4A). The difference between the time at which 50% of the max density is reached for the uninduced cells and the induced cells, Δt50, is used as a metric that positively correlates with aggregate size. A detailed overview of this assay is shown in S10.

**Figure 4.**
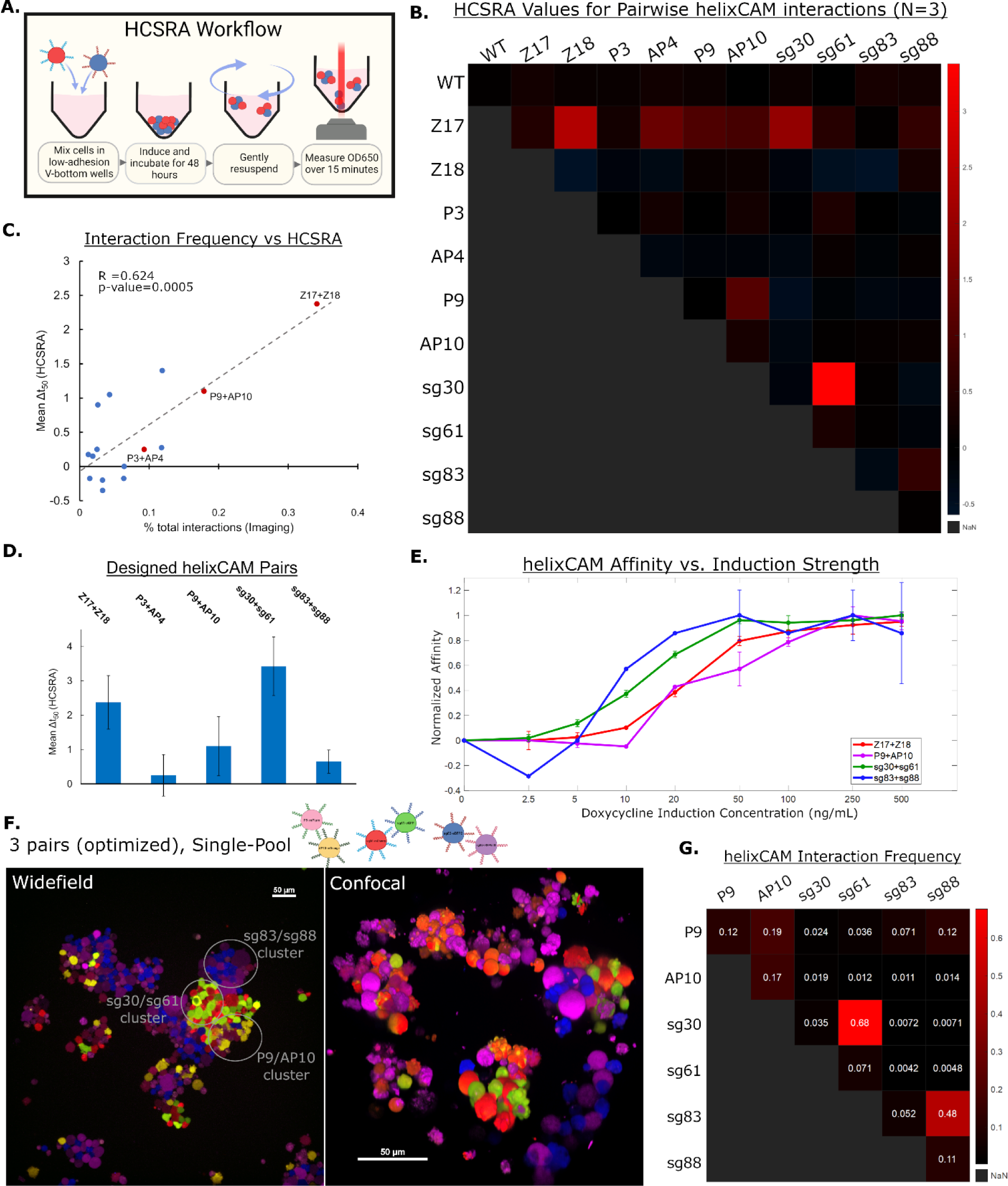
HCSRA measurement of helixCAM affinity and optimized three-pair aggregation. **A.** Cartoon of Human Cell Sedimentation Rate Assay (HCSRA). **B.** Heatmap showing Δt_50_ values from HCSRA for helixCAM affinity for self, pairwise, and wild-type conditions. Each square represents the mean of three replicates. **C.** Comparison of Δt_50_ values from HCSRA to pairwise interaction frequencies from Figure 2D. A linear correlation was observed, with an R_2_ of 0.624 and p-value of 0.0005, using the F-statistic. **D.** Bar graph of Δt_50_ for complementary helixCAM pairs, demonstrating a range of affinity. Error bars represent S.D., N=3. **E.** Dose curve of helixCAM-induced affinity for the top four helixCAM pairs. Δt_50_ for each pair is normalized to each pair’s maximum value. Error bars represent S.D., N=2. **F.** P9^mPlum^, AP10^mOrange^, sg30^mCherry^, sg61^eGFP^, sg83^iRFP670^, and sg88^eBFP2^ cells induced to express helixCAM within a single mixed culture. This subset of helixCAMs was selected as the most orthogonal set from HCSRA results. Clear sub-clusters can be observed. Widefield image was taken at 20X with a 5x5 tile across five channels and cropped to show regions of interest (Uncropped image in S12). Confocal image was taken at 40X magnification with a spinning disc confocal system as a Z-stack and reconstructed using Nikon Elements. **G.** Interaction frequency table derived from the optimized three-pair co-culture image. Designed colocalization can be observed in all three helixCAM pairs. A detailed description of the analysis pipeline can be found in S5.

Using the HCSRA, the Δt50 for all self and pairwise helixCAM interactions were measured, along with interaction with wild-type K562 cells (Figure 4B). It was noted that the Δt50 values for interactions between the original six helixCAMs correlated with interaction frequencies derived from microscopy (Figure 4C), with an R^2^ of 0.624 and a linear model p-value of 0.0005. The agreement of the HCSRA Δt50 values to the microscopy-derived interaction frequency supports the intended function of HCSRA as a readout for the affinity between cells. Additionally, we observed that, in agreement with microscopy observations, both library-derived helixCAM pairs demonstrated high affinity and specificity. Indeed, the sg30/sg61 pair surpasses the binding affinity of the previously highest-affinity pair, Z17/Z18 (Figure 4D).

For the four highest-affinity helixCAM pairs, we investigated whether the magnitude of the aggregative effect can be adjusted by changing helixCAM expression levels. Affinity across a range of induction strength was tested (Figure 4E), revealing that helixCAM cell pairs could be tuned to form aggregates of varying sizes solely by controlling helixCAM expression. These findings suggest that helixCAM-induced affinity could be easily calibrated for individual applications through controlling helixCAM expression with small-molecule inducers.

Using the most orthogonal set of helixCAM pairs (P9/AP10, sg30/sg61, sg83/sg88), we repeated the three-pair co-culture experiment. Similar to the previous set, striking segregation was observed in cell aggregates (Figure 4F). In this case, quantification of interaction frequencies revealed that the new set had notably fewer inter-pair interactions, leading to frequent complementary interactions and few off-target interactions (Figure 4G, S12). These results show that, by utilizing the HCSRA results, it is possible to select multiple compatible helixCAM for directing parallel cell adhesion events within a single culture.

### helixCAMs enable additive construction of sophisticated three-dimensional cell structures

Specific spatial patterning of different specialized cell types is a hallmark of human tissue, where the organization enables functions such as nutrient exchange and cell signaling [42]. We hypothesized that, by using a “core” cell expressing a single helixCAM followed by sequential incubation with “layer” cells expressing two helixCAMs, it would be possible to replicate complex layers of cells, similar to those found in human tissues. To this end, we selected the sg30^mCherry^ cell as the “core” cell and created intermediate three “layer” cell types, each expressing two orthogonal helixCAMs (sg61+sg88^eBFP2^, SG83+P9^iRFP670^, P10+Z18^eGFP^).

The goal was to form stable inner layers using the most orthogonal helixCAMs while relegating the more promiscuous Z17 for the outer layer (Z17^eCFP^) (Figure 5A).

**Figure 5.**
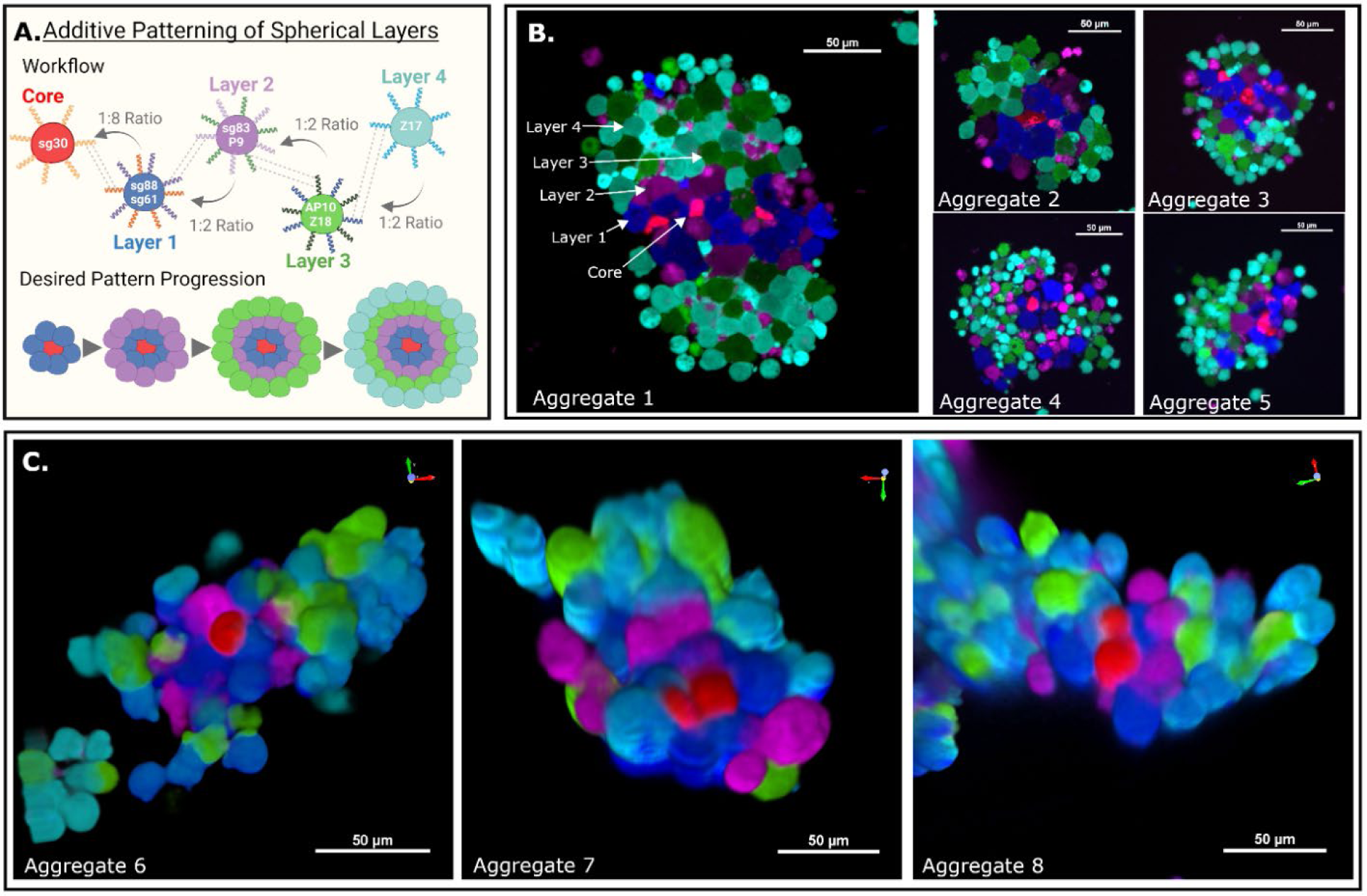
helixCAMs enable additive construction of sophisticated three-dimensional cell structures. **A.** Schematic of spherical layering workflow. Five K562 lines were built, expressing either one or two helixCAMs and an identifying fluorescent protein. Layer 1 cells were mixed with core cells at an 8:1 ratio, and subsequent layers were then mixed at a 2:1 ratio. **B.** Five select aggregates resembling the desired patterning are shown. Arrows overlaid on Aggregate 1 indicate the location of each cell type. Aggregates were imaged at 20X magnification and cropped for emphasis. **C.** Confocal imaging of patterned spherical structures. Core and layer cells are visible and near intended locations. Confocal images were taken at 40X magnification with a spinning disc confocal system as a Z-stack and reconstructed using Nikon Elements.

A number of aggregates formed through this process demonstrate closely resemble the envisioned structural layout (Figure 5B and 5C). Unsurprisingly, the most consistent interactions were the SG30/SG61-induced interactions between sg30^mCherry^ “cores” and sg61+sg88^eBFP2^ “layer 1” cells. We also observed a layer of magenta SG83+P9^iRFP670^ “layer 2” cells coating the blue “layer 1 cells,” induced by SG83/SG88 interactions. The next expected layer would be the P10+Z18^eGFP^ “layer 3” cells, which are present to some degree. However, a significant number of green “layer 3” cells were also found intermixed with Z17^eCFP^ “layer 4” cells rather than forming clearly delineated layers, likely a consequence of the high affinity of the Z17/Z18 interaction. Additionally, we observed that, whereas initial layers formed spherical structures, unevenness in these seed aggregates appear to be amplified by subsequent cell layers, often leading to cells from layers 2-4 organizing on opposite sides of the core and forming a hamburger-like presentation. The striking complexity formed with four helixCAM pairs, and the resemblance of aggregates to the intended multicellular structure blueprint, alludes to the potential for helixCAMs to guide the patterning of sophisticated multicellular structures.

### helixCAMs enable targeting of suspension cells to adherent cells

In addition to tissue construction, cell-cell interactions are critical to the targeting of immune cells to areas of active infection or malignancy[43], [44]. We hypothesized that, if adherent cells of interest can be made to express one member of a helixCAM pair, it would be possible to recruit suspension cells expressing a complimentary helixCAM to that region. To start, we generated adherent HEK293 cell lines capable of expressing helixCAMs. After allowing them to grow to confluency, we added corresponding K562 helixCAM-expressing cells, then washed off unbound cells. While few suspension cells remained bound to the adherent wild-type and uninduced helixCAM cells, copious suspension cells were bound to the induced adherent helixCAM cells throughout multiple washes (S13). A similar binding experiment was performed between adherent HEK293 helixCAM cells and a trypsinized suspension of complementary HEK293 helixCAM cells (Figure 6A). As with K562 cells, strong helixCAM- dependent enrichment of bound suspension HEK293 cells was observed, and, by 24 hours, the bound cells had re-established their adherent morphology (Figure 6B). These experiments indicate that helixCAM is a viable tool for targeting suspension cells and adherent cells alike to pre-established adherent cells of interest.

**Figure 6.**
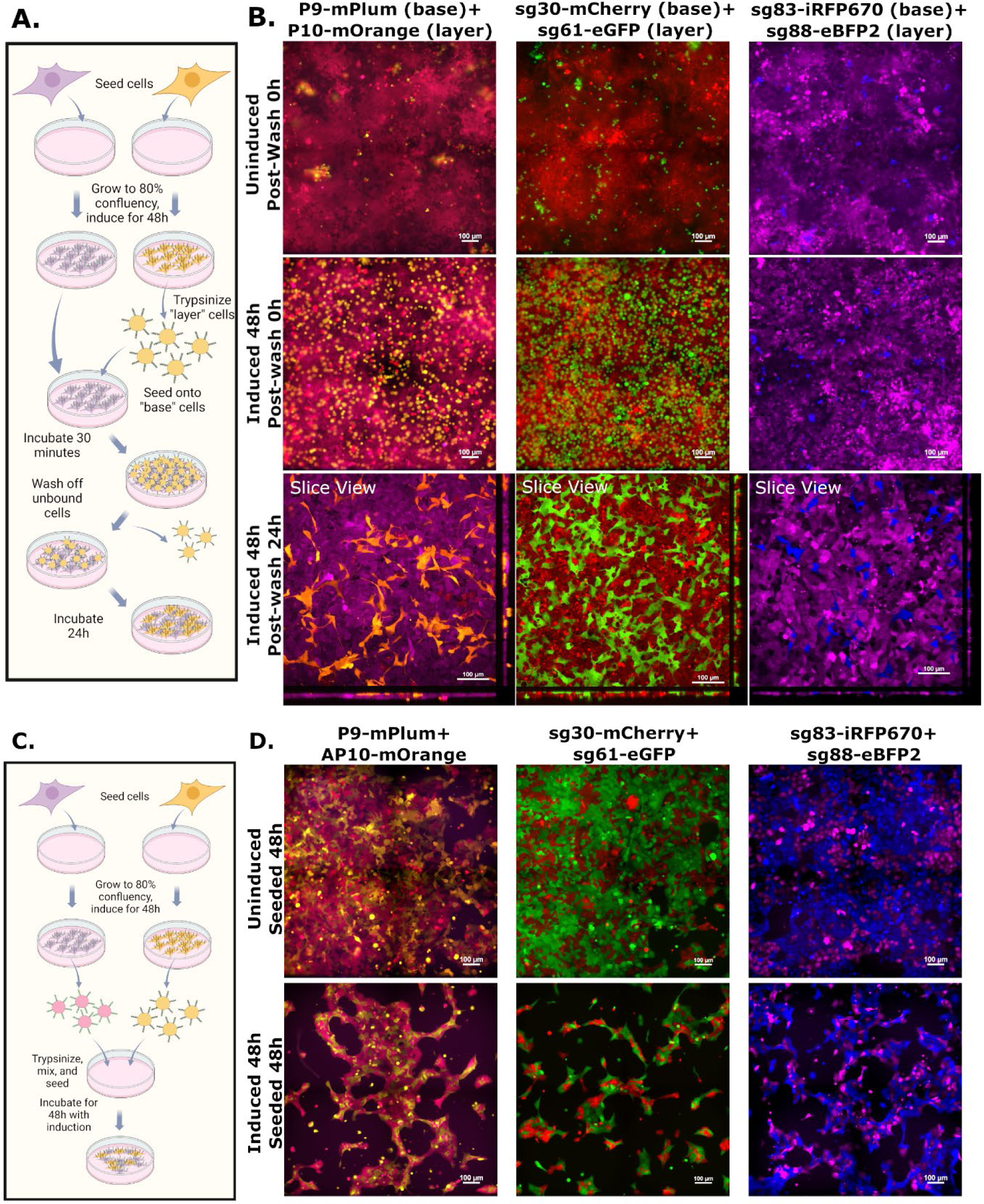
helixCAMs enable targeting of suspension cells to adherent cells and can impact adherent cell morphology. **A.** Workflow for targeting suspended HEK293 cells to adherent HEK293 cells. After cells were grown to confluency, one population is trypsinized and added to the other population, after which unbound cells are washed off. Bound cells are then allowed to re-establish an adherent morphology. **B.** Panel images show three helixCAM pairs uninduced, induced 0h, or induced 24h. Uninduced and induced 0h conditions were imaged using a widefield microscope at 20X magnification with 2x2 tiling, and induced 24h images were taken using a confocal microscope as a Z-stack, also at 20X magnification with 2x2 tiling, shown as “slice view.” **C.** Workflow for joint seeding of complementary helixCAM-expressing HEK293 cells. Cells were seeded and induced, followed by trypsinization and re-seeding as a single mixture. **D.** Panels show the distribution and morphology of co-cultured helixCAM HEK293 cells for three helixCAM pairs. Uninduced cells establish normal cell morphology, whereas induced cells form long, stretched bundles.

### helixCAM interactions impact adherent cell migration and morphology

Adherent cells, such as HEK293s, express various adhesion proteins that mediate strong interactions with the plate surface and with each other. To examine whether the strength of helixCAM interactions was capable of competing with endogenous adherence, pairs of HEK293 helixCAM cell lines were co-cultured and examined for preferential interactions by widefield microscopy (Figure 6C). While the two cell types in uninduced controls were evenly distributed and displayed normal cell morphology, induced cells were much more locally concentrated, left larger areas of the well empty, and formed elongated bundles (Figure 6D). Within each bundle, it is possible to identify the cell bodies through alternating fluorescence, and rather than the rotund morphology found in the uninduced conditions, the induced cells appear stretched, with some cell bodies spanning over 100μm (uninduced cell lengths were closer to 30μm). The notable change in morphology from HEK293 helixCAM cells indicates that helixCAM interactions can compete with endogenous interactions to influence adherent cells’ migration and morphology.

### helixCAMs enable tunable and simultaneous patterning of multiple cells types on CC- patterned surfaces

While the experiments above focus on helixCAMs’ capabilities to bind cells to one another, the affinity and specificity of helixCAMs could also be leveraged to pattern multiple cell types onto a surface, enabling purification of cells from a mixed population or patterning of microfluidic devices. To test helixCAM’s capacity to pattern cells, we expressed and purified both His-tagged CC peptides as well as His-tagged GFP-CC fusion proteins for CCs sg30, sg61, sg83, and sg88 and confirmed the quality of the purified protein species with LC-MS (S14). The protein solutions were applied to nickel-coated plates to create the CC-coated surface, followed by the addition of helixCAM-expressing cells for patterning. After removing unbound cells, the remaining cells were then imaged and quantified (Figure 7A).

**Figure 7.**
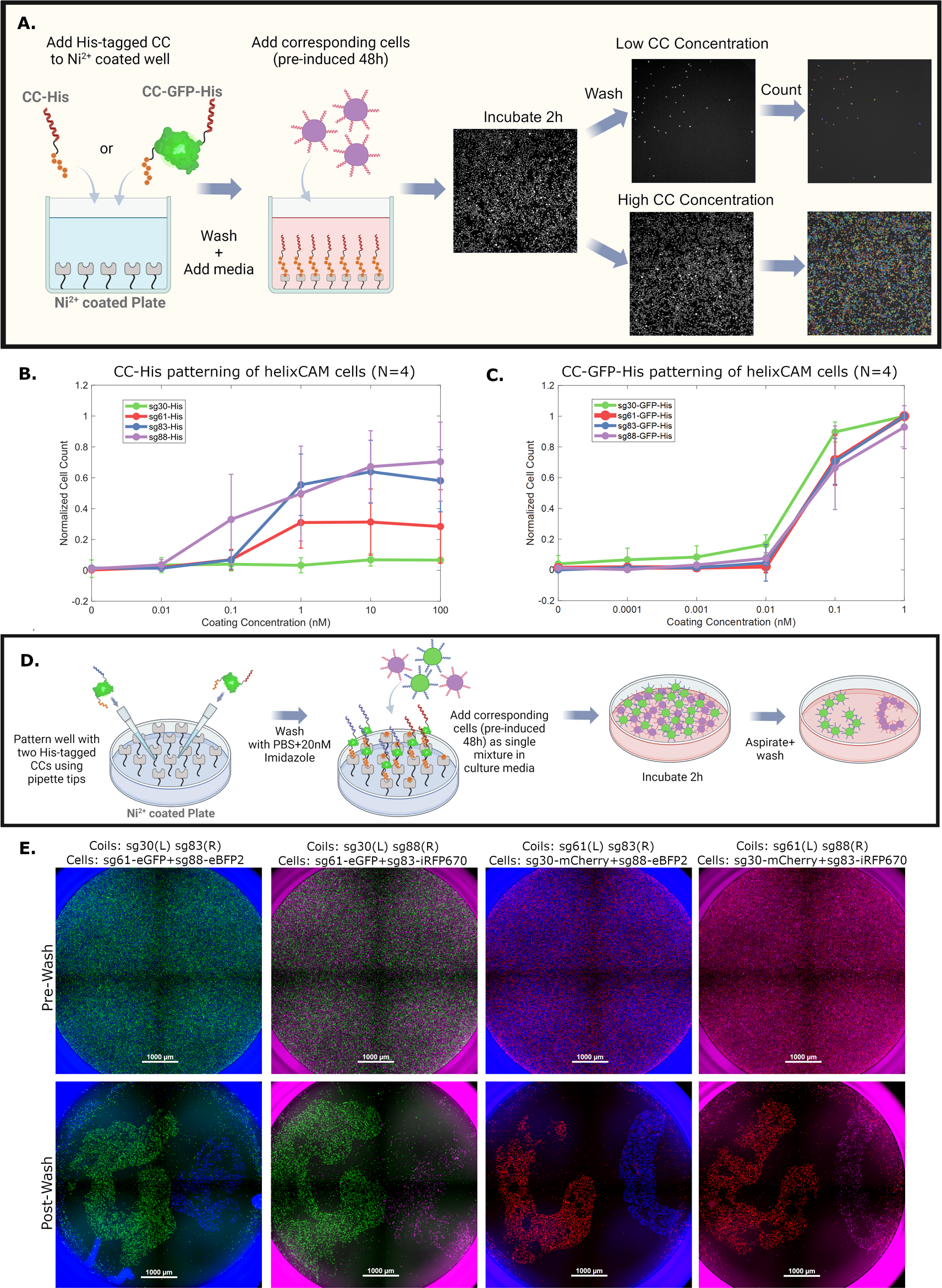
helixCAMs enable tunable and simultaneous patterning of multiple cells types onto CC-patterned surfaces. **A.** Workflow for CC surface patterning across a gradient and downstream automated image segmentation and analysis. Wells are coated with His-tagged CCs or His-tagged CC-GFP fusion protein solutions, then cells are added. Unbound cells were washed off, and the entire well was imaged at 4X magnification as a 2x2 tile. Images were segmented and cells counted by CellProfiler. **B.** Normalized bound cell count is plotted against CC coating concentration for each of four CC-His peptides tested. Bound cell counts were normalized to the maximum in each channel to reduce variance from segmentation (raw counts in S15). Reported values are medians with N=4 and error bars are S.D. **C.** Normalized bound cell count is plotted against CC coating concentration for each of four CC-GFP-His peptides tested. All four proteins demonstrated strong capability for binding complementary helixCAM cells. Reported values are medians with N=4 and error bars are S.D. **D.** Workflow for simultaneous patterning of two cell types within one well. Two distinct CC-GFP-His protein solutions were added in a “G” or a “C” pattern. The two complementary helixCAM cell populations are added as a single mixture to cover the well surface. Unbound cells were then washed off and the bound cells imaged at 4X magnification as a 2x2 tile. **E.** Dual-CC-patterned wells with two cell populations pre- and post-wash. Pre-wash, cell populations are evenly distributed and fully cover the plate surface. Post-wash, helixCAM cells complementary to the CC patterned at the G or C locations remain bound, but the non-complementary cells are not bound to those locations, and outside of patterned regions, few cells are observed.

Using this method, we tested five orders of magnitude of protein concentrations and an untreated condition for all eight CC fusions. Results from four replicates were normalized by channel to control for variability due to image segmentation, and the median and standard deviation reported in Figure 7B for CC-His coating and Figure 7C for CC-GFP-His (raw counts can be found in S15). For all conditions but sg30-His, the CC-coated surface led to helixCAM cell binding, and the number of cells bound strongly correlated to the concentration of protein coating applied. Notably, at the same coating concentration, CC-GFP-His-coated wells bound markedly more cells than for CC-His-coated wells, and, at higher concentrations, led to cells covering the entirety of the plate surface.

Next, we tested whether it would be possible to simultaneously bind two distinct helixCAM cell populations to the plate in a specified spatial pattern. To test this, we pre-patterned each well with two distinct CC-GFP-His proteins, either in a “G” shape on the left or a “C” shape on the right, added a mixture of the two corresponding K562 helixCAM cells, then removed unbound cells (Figure 7E). With four CC options, there are four compatible 2-CC combinations: sg30+sg83, sg30+sg88, sg61+sg83, and sg61+sg8. Combinations between complementary CCs, such as sg30+sg61, are not compatible for this purpose, as the corresponding helixCAM cells would form aggregates in suspension. Figure 7F shows the pre- and post- washed CC-GFP-His-patterned, helixCAM cell-coated plates. The pre-wash images exhibit uniform distribution of the two helixCAM cell types across the well surface. After washing, dense helixCAM cells can be found localized to the corresponding “G” and “C” CC- patterned regions, whereas untreated regions are almost entirely free of cells. The same experiment, using dots instead of letters as patterns, replicates this result (S16). The strong and programmable binding of helixCAM cells to CC-patterned surfaces demonstrates its potential as a method for highly specific cellular pull-down and patterning, with the number of simultaneous cell types patterned determined by the number of orthogonal CC-pairs available.

## Discussion

In this study, we presented helixCAM, an extensible platform for engineering programmable cell-cell interactions in both bacteria and human cells. helixCAMs were demonstrated to induce formation of large, patterned cell aggregates containing thousands of cells and spanning multiple cell layers. From helixCAMs composed of previously established CC domains, we rationally designed CC peptide libraries and, after screening for affinity and specificity, found two novel CC pairs with favorable helixCAM properties. We then tested high-affinity helixCAM pairs across a range of cell types and applications, demonstrating their versatility in guiding multicellular patterns and morphologies.

One of the most complex applications tested was the construction of sequentially layered spherical aggregates using five engineered cell types (Figure 5), from which a number of interesting observations arose. Based on pairwise affinity data, we expected the Z17-eCFP cells to be the least predictable due to their promiscuity and placed them last in the coating sequence to mitigate its impact. Interestingly, the strength of the Z17/Z18 interaction appeared to overpowered previously formed P9/AP10 interactions, pulling the green “layer 3” cells away from “layer 2” cells to form sub-clusters rather than clearly delineated layers. This observation is consistent with the “differential adhesion hypothesis” of how cells in an aggregate naturally self-segregate based on binding affinity – a key element of biological development[45]. The potential for subsequent helixCAM interactions to disrupt previously-formed ones indicates a need for independent tuning of the expression level of each helixCAM. It may also be possible to guard against disruption by subsequent cells by solidifying interactions between additions with an intermediary step of homodimer CAM expression, such as cadherins. Relatedly, the oval structures that formed instead of the desired spherical structure may be due to a combination of the initial core-layer1 aggregates not forming perfectly spherical structures, and the strength of subsequent layer interactions accentuating the imperfections.

We envision several potential biological applications enabled by the helixCAM in immunology. For instance, intercellular signaling is known to be critical to the activation of immune cells[43], [44], but few synthetic methods for directing or augmenting cell-cell interactions exist beyond the use of endogenous CAMs. By using helixCAMs, small molecule inducers could be used to adjust the binding strength of immune cells to their targets in conjunction with B- or T-cell receptors and other co-stimulatory signals, leading to a more refined understanding of the physical role of cell interactions in immune cell activation. In addition, helixCAMs can augment weaker activation events, such as those of chimeric antigen receptors, either through constitutive expression or triggered to be expressed by other binding events. helixCAMs may even be used to directly localize immune cells to infected or malignant tissues, given that those cells can be transduced to express helixCAMs or functionalized with targeting CCs (for instance, with a CC-antibody fusion).

Tissue engineers may also find compelling applications for helixCAMs. Current paradigms for engineering tissues for therapeutic purposes, such as 3D cell printing[9]–[12], are constrained by printing resolution and limited survival of printed cells [13], [14]. As demonstrated in Figure 5B, helixCAMs can be used to manufacture complex spherical structures with single-cell level accuracy. By expressing helixCAMs in physiologically-relevant cells, it may be possible to create smaller high-resolution building blocks that are subsequently arranged into more complex tissues[46]. Furthermore, helixCAMs can be integrated naturally into existing tissue engineering workflows. For instance, for platforms that grow[47] or 3D print vasculature[48], by utilizing helixCAM, it should be possible to functionalize the synthetic vasculature with additional cell types to enhance its resemblance to physiological vasculature.

The helixCAM platform’s versatility extends beyond cell-cell interactions to cell-surface interactions, opening the door to opportunities to augment conventional cell purification and patterning methods. For instance, by expressing helixCAMs in the desired cell population, those cells can be isolated using CC-coated magnetic beads. Another area benefiting from programmable cell-surface interactions is organ-on-a-chip platforms [49]–[51], where it may be desirable to culture multiple cell types in specific spatial orientation. By pre-patterning the microfluidic device with multiple CC species, complex cell distribution patterns could be designed to further mimic the physiological environment.

The helixCAM platform offers a simple but powerful way to control cell-cell interactions, a ubiquitous feature found from prokaryotes to eukaryotes. Beyond the set of five helixCAM pairs characterized in this work, many more orthogonal helixCAMs likely can be discovered and further expand potential applications. We believe that the toolkit of helixCAMs presented here will assist biologists across multiple fields to better understand the roles cell interactions play in various contexts, as well as enable engineers to better control these interactions for therapeutic applications.

## Acknowledgement

We are extremely grateful to Dr. Roman Jerala for sharing the engineered coiled-coil pairs P3/AP4 and P9/AP10. We are grateful to Dr. Bridget Baumgartner, Dr. Justin Gallivan, Dr. Jesse Dill, and Dr. Joseph Pomerening for helping to fund and shape the direction of our work. Thank you to Dr. Ron Raines at MIT for the use of LC-MS equipment for characterizing the CC-His and CC-GFP-His proteins. We thank Dr. Paula Montero Llopis and Ryan Stephansky from the HMS MicRoN core facility for their assistance in microscopy techniques and maintenance of microscopes. We also thank the HMS Immunology Flow Cytometry facility, particularly Chad Araneo, Jeff Nelson, and Meegan Sleeper for training and maintenance of the FACS machines. Thank you to Songlei Liu for providing the v-bottom ULA plates for HCSRA experiments. We are extremely grateful to Tiffany Dill and John Aach help to edit the manuscript. Thank you to Emma Taddeo for all of her help with the lab’s administrative work, and to Nicole D’Aleo for managing the grants. Plasmid design and map image generation was done in A Plasmid Editor (ApE). A majority of schematic figures made in this work were created with BioRender.

This work was supported in large part by the DARPA Engineered Living Materials program under contract W911NF-17-2-0079. G.C. was also supported by the NHGRI Centers of Excellence in Genomic Science (RM1HG008525) as well as a general gift from the Zhijun Yang Research Fund. T.W. was also supported by the DoE under the award DE-FG02-02ER63445.

## Author Contributions

Conceptualization, G.C., T.W., E.A., and G.M.C.; Methodology, G.C., and T.W.; Investigation, G.C., C.T., N.B., A.C., T.L, T.W., and E.A.; Analysis/Software, G.C., N.B., C.T., and A.C.;

Writing – Original Draft, G.C.; Writing – Review & Editing, G.C., A.C., C.T., T.W., and T.L.; Visualization, G.C., A.C., C.T., and T.W.; Supervision, G.C., T.W., and G.M.C., Funding

Acquisition, G.W., T.W., E.A., and G.M.C., Resources, G.M.C

## STAR Methods

### RESOURCE AVAILABILITY

#### Lead Contact

- Further information and requests for resources and reagents should be directed to and will be fulfilled by the lead contact, George Church.

#### Material availability

- Plasmids for constructing helixCAM lines have been deposited to Addgene.

#### Data and code availability

- All original code has been deposited at Zenodo and is publicly available as of the date of publication. DOIs are listed in the key resources table.
- Any additional information required to reanalyze the data reported in this paper is available from the lead contact upon request.

### EXPERIMENTAL MODELS AND SUBJECT DETAILS

#### E. coli

Standard cloning strains DH5a and DH10B were used for most cloning and DNA preparation purposes. For bacterial adhesion assays, the T7 Express Competent *E. coli* cells were used (NEB C2566). Libraries were transformed using E. cloni® 10G Supreme Electrocompetent Bacterial Cells (Lucigen 60107-1). CC-His proteins were expressed in BL21(DE3) cells (Novagen 69450). All cultures were grown in LB Miller media at 37°C.

#### S. Cerevisiae

Yeast strains for each haploid yeast strain (MATa/ɑ) were gifted by David Younger and the Klavins Lab. Yeast strains were grown at 30°C in YPD or Yeast SDO media. **Cell Lines:** K562 cells (ATCC CCL-243) were used for most human helixCAM line generation. They were cultured in either shaking or standing suspension and passaged at a concentration of 2x10^6^/mL to a seed concentration of 2x10^5^/mL. Adherent cells used for targeting experiments were the AAV-293 cells (Agilent 240073) for their superior adhesion. They were seeded at 10% confluency and allowed to grow to 90% confluency before passaging. All cells were cultured at 37°C and 5% CO^2^ in DMEM+GlutaMax (ThermoFisher 10566016) supplemented with 10% Fetal Bovine Serum (Corning 35-010-CV) and 1% Penicillin-Streptomycin (ThermoFisher 15070063)

### METHOD DETAILS

#### Plasmid Design and Construction

For *E. coli* helixCAM construction, existing coiled-coil sequences were found via literature search and private communications and synthesized (Twist). Coils-encoding DNA fragments were cloned into a pQE protein expression vector (Qiagen pQE-80L) with in-frame with a flanking leader sequence and an *EhaA* coding sequence via Gibson Assembly (NEB E2611L). Plasmids were transformed in T7 Express Competent *E. coli* cells (NEB C2566), miniprepped (Qiagen 27104), and verified via Sanger sequencing (Genewiz).

To construct the coiled-coil-sGFP libraries, DNA fragments containing the full sGFP expression cassette with flanking Golden Gate Assembly overhangs were synthesized (IDT). The plasmids were then cloned into the pET-DEST T7 expression vector (ThermoFisher 12276010), creating a final single plasmid containing all three subunits of GFP (S1). Coiled-coil sequences were PCR-amplified from gene fragments using Q5 Polymerase (NEB M0492L) with *BsaI* recognition sites and unique 4 base pair overhangs to either fuse the fragment 3’ of GFP10 or 5’ of GFP11 using Golden Gate Assembly (NEB E1601L). Assembled plasmids were then transformed into E. cloni® 10G Supreme Electrocompetent Bacterial Cells (Lucigen 60107) Cloned plasmids were miniprepped and transformed into T7 Express Competent E. coli strain (NEB C2566) for expression and screening.

To construct coiled-coil libraries for yeast SynAg screening, the SynAg plasmid vectors were used (gifted from the Klavin Lab). The carbenicillin resistance cassette was replaced with kanamycin resistance to prevent pQE plasmid contamination. Additionally, silent mutations were made to the *SUMO* coding region in the SynAg-a cassette via site-directed-mutagenesis (NEB E0554S) to distinguish from SynAg-alpha during amplification. Coiled-coil sequences were amplified from the previously-described *E. coli* pQE vectors. Correct amplification was determined through individual qPCR reactions and representative PCR products were checked via gel electrophoresis. PCR products were pooled and assembled into the SynAg vectors via Gibson Assembly. Assembled plasmids were purified and electroporated into E. cloni® 10G Supreme Electrocompetent Bacterial Cells (Lucigen 60107). Cloning efficiency was quantified via plating of dilutions onto LB+Kanamycin and counting colonies. Cultures were grown with selection overnight and then miniprepped to produce plasmid DNA for transformations. Library scale lithium acetate transformations of *S. cerevisiae* were performed as described in Younger *et al.* [41], [52], with the adjustment of 10μg of PmeI digested plasmid DNA and fresh salmon sperm DNA. Dilutions onto selective yeast Synthetic Drop-Out -trp agarose plates were performed, and transformation efficiency was confirmed to be >300-fold coverage of the coiled-coil library size.

To construct plasmids for human helixCAM experiments, the Thermo pDisplay was used (ThermoFisher V66920). Coiled-coil sequences were flanked with Gibson Assembly overhangs and inserted in-frame between the IgK leader peptide and *PDGFR* sequences through Gibson Assembly (NEB E2611L). The full-length human helixCAM construct was amplified through PCR and inserted behind the pTet promoter of the PiggyBac Tet-On vector (Takara 631168). An identifying fluorescent protein sequence with a T2A sequence was placed between the T2A sequence and the puromycin resistance cassette of the Tet-On vector. For the blasticidin-resistance plasmid (for dual-helixCAMs), the fluorescent protein sequence and the puromycin resistance gene were replaced with a blasticidin resistance gene, which was cloned from Addgene Plasmid 74918 (gifted from Jose Silva Lab).

Plasmids for *E. coli* expression of His-tagged coiled-coil peptides was done by first synthesizing codon-optimized gBlocks (IDT) comprising each Coiled-Coil with a C-terminal 6xHis tag. The DNA fragments were inserted into a pET22b expression vector (Novagen 69744) via Gibson assembly (New England Biolabs E2611S). Plasmids for expression of His-tagged CC-GFP fusion protein were done by replacing the CMV promoter of the pcDNA3.1 vector (ThermoFisher V79020) with the CAG promoter, followed by insertion of the coiled-coil gBlock (codon-optimized for mammalian expression) N-terminally to a TEV cleavage sequence, GFP, and 6xHis fusion (S1).

#### helixCAM K562 Cell Line Construction

PiggyBac-flanked donor plasmids were packaged with the Super PiggyBac transposase vector (SBI PB210PA-1) using Lipofectamine 2000 (ThermoFisher 11668019) at a 1:1 ratio in Opti-MEM (ThermoFisher 31985062). For K562 cells, the liposome-DNA solution was first added to a well, then K562 cells were added in culture media on top. For HEK293 cells, cells were allowed to first grow to 70% confluency, then the liposome-DNA solution was added dropwise to the media. 48 hours after transfection, cells were changed into selection media containing either puromycin (ThermoFisher A1113802) or blasticidin (ThermoFisher A1113903). For K562 cells, the selection concentration used was 1μg/mL for puromycin and 6 μg/mL for blasticidin, and for HEK293, it was 2μg/mL for puromycin and 10 ug/mL for blasticidin. After four days of selection, cells were passaged to recover for 48 hours, then subsequently single-cell sorted (BD FACSAria), either with fluorescence selection (for puromycin lines) or only through FSC/SSC discrimination (for blasticidin lines). Subsequently, surviving lines were imaged and scored for fluorescence and viability, and the ones scoring highly in both were selected.

#### Pairwise helixCAM Aggregate Formation and Sedimentation Assays

helixCAM interactions in *E. coli* were tested as follows. A single colony was picked into 3mL of LB+carbenicillin, and grown overnight. 150uL of culture was diluted into 30mL LB+carbenicillin and grown for 75 minutes. 60mg of arabinose was added to induce, and culture was grown for an additional 3 hours. 2mL from two separate cultures, each expressing a distinct helixCAM, were mixed, and after four hours, a pipette is placed a quarter of the depth into the culture, and 100uL of cells are taken. This was either measured as an OD600 or plated onto a #1.5 glass coverslip for imaging.

helixCAM interactions in K562 cells were tested as follows. 2x10^5^ of two distinct K562 helixCAM cell lines were co-cultured in a low-adherence 24-well plate (Corning 3473) in culture media with 500μg/mL doxycycline added. Cells were incubated for 48 hours, then either mounted using a #1.5 glass coverslip for widefield imaging or embedded in low melting point agarose (ThermoFisher 16520050) for confocal imaging using wide-bore pipette tips (Rainin 30389191). For multi-pair co-cultures, the same process, including the number of cells of each line, was followed.

For the Human Cell Sedimentation Rate Assay (HCSRA), 1.75x10^5^ of each cell line to be tested (or 3.5x10^5^ for self-interaction) was added to 150uL of culture media with or without doxycycline. The cells were cultured in a specific ultra-low-adhesion v-bottom plate (SBio MS- 9096VZ), which has the right v-bottom curvature allowing this assay to work. The cells were allowed to interact for 48 hours on an orbital shaker at 150rpm. To measure, the cells were resuspended using 15 seconds of shaking at 900rpm, then immediately placed into a plate reader (Molecular Devices M5) on the same bench, and kinetic OD650 was measured over 15 minutes. The data is then exported and analyzed in MATLAB.

#### Coiled-Coil Library Screening

For stage one of the CC library screen, a tripartite split-GFP method was used. Colonies were picked from strains containing each paired library combination and grown in 3mL LB+Carbenicillin (50 ng/μl) overnight. The culture was diluted 1:100 in 100μL of LB carbenicillin and grown in shaking culture in a BioTek Synergy H1 plate at 567 cycles per minute and 37°C for 1hr. Next, the cells were induced at 1:1000 with IPTG (Teknova I3431) and grown for 45 minutes. Cells were then diluted 1:100 into PBS (ThermoFisher 10010023) and sorted on a BioRad S3e Cell Sorter. From each library, cells were sorted into for the top 0.5% GFP intensity, and the sorted cells were grown to a high density and sorted again, selecting for the top 2% of GFP intensity. Plasmids were miniprepped from the resulting population and sequenced through NGS. 102 coiled-coil candidates demonstrating the highest affinity and orthogonality were selected for the next stage of the screen.

The next stage of the screen used the yeast SynAg assay. Haploid MATa and MATɑ strains expressing the SynAg cassette were picked from fresh colonies into 3mL of YPD and grown in a drum rotator for 24 hours. 2.5μL of MATa and 5μL MATɑ saturated cultures were added to a fresh 3mL of YPD and grown for 24 hours. For optimization of the SynAg system, cells were diluted 1:200 in 1X PBS for assay through flow cytometry (Miltenyi MACSQuant VYB) of the expression of mCherry and mTurquoise. For screening CC candidates, diploid selection was done through complementary lysine and leucine auxotrophic markers, then enriched for an additional 24 hours before genomic DNA preparation and NGS. We noticed that the original aga1p/aga2p yeast display system [53] led to low surface presentation of the CCs, which we attributed to protein degradation. By inserting a SUMO-tag [54] between the coiled-coil and the ga2p domain, the membrane protein was stabilized, which increased surface presentation (S8) and led to successful yeast mating events.

#### Next-Generation Sequencing of screened CC candidates

Plasmid DNA was extracted from each library and amplified via PCR (NEB M0492L) with primers containing homology to the binding region of the indexing primers. PCR reactions were monitored via qPCR (SigmaMillipore S4438) on a Roche LightCycler 96. Indexing of the library member DNA proceeded with a global primer (Pi5) and a unique primer (Pi7) for sample barcoding. Sequencing was performed at the Biopolymers Facility at Harvard Medical School on an Illumina NextSeq 500 sequencer, yielding 300bp forward and reverse reads containing both coils when combined.

#### helixCAM Additive Spherical Patterning

To build the spherical structure, all five cell types (core and layers 1-4) were pre-induced for 48 hours with 500ng/mL doxycycline in culture media. Core cells were added to layer 1 cells in a 1:8 ratio, followed by a 12-hour incubation. After incubation, aggregates were resuspended through gentle vortexing, and a pellet was allowed to form before removing the unbound cell supernatant. Subsequent cell populations are added at a 1:2 ratio, following the same steps. After the last 12-hour incubation and unbound cell removal, aggregates were diluted in culture media and either mounted using a #1.5 glass coverslip for widefield imaging or embedded in low melting point agarose (ThermoFisher 16520050) for confocal imaging.

#### HEK293-HEK293 and HEK293-K562 Cell Targeting

Complementary helixCAM lines of either two HEK293 cell types or one HEK293 and one K562 are grown to 80% confluency on a 24-well glass-bottom plate (Mattek P24G-1.5-10- F). Doxycycline is then added at 500ng/mL and cells were induced to express helixCAMs for 72 hours. In the case of HEK293-HEK293 targeting, one population of HEK293 cells were trypsinized to be temporarily suspension cells, then added to the second still-adherent population of helixCAM-expressing HEK293 cells. For HEK293-K562 targeting, the K562 cells were directly added to the adherent HEK293 cells. The coated cells were allowed to incubate for 30 minutes then imaged (pre-wash). Unbound cells were then aspirated and washed with culture media+doxycycline twice. The cells were then imaged again (post-wash). For HEK293- HEK293 targeting, cells were returned to the incubator for an additional 24 hours of growth before imaging again to observe the re-establishment of adherent cell morphology.

#### Expression and Purification of His-tagged CC and His-tagged CC-GFP

To express His-tagged CCs, *E. coli* cells (BL21 DE3) were grown in Terrific Broth (TB) overnight at 25°C with 200 μg/mL ampicillin and glucose to 0.5% till they reached an OD of ∼0.5. They were then induced with 1mM Isopropyl ß-D-1-thiogalactopyranoside (IPTG) and grown overnight at 16°C. The cultures were harvested via centrifugation at 6000xg for 15min at 4°C, pellets resuspended in 20mM Tris, 500mM NaCl, 20mM Imidazole, 500μM PMSF at a pH of 8.0, then lysed via sonication. The resulting lysate was then spun at 30,000xg at 4°C for 90 minutes to remove insoluble proteins and cell debris. The supernatant was passed through a 0.45μm filter then run on a HisTrap HP 1mL column (Cytiva 17-5247-01) and fractions were collected using an Akta Pure FPLC. The fractions were run on an SDS-PAGE gel (BioRad 4569033) and stained via Coomassie (BioRad 1610786) to select fractions with significant protein. Pooled fractions were then dialyzed using Slide-a-Lyzer mini dialysis cassettes with a 3.5k MWCO (Thermo Fischer, Cat#88403) into 50mM Tris, 100mM NaCl, pH 7.6. Some contaminating proteins precipitated during dialysis so the solutions were clarified by spinning at 15,000xg for 15min at 4°C. Yields ranged from 6-25 mg of protein per liter of culture.

To express His-tagged CC-GFP fusion protein, HEK293T cells were cultured to 70% confluency in a T75 flask, then transfected with 15μg of CC-GFP-His expression plasmid using PEI. Cells were grown for 72 hours, then lysed in 2mL of Triton buffer (50mM Tri pH8, 150mM NaCl, 0.1% Triton X-100). His-tagged proteins were bound to Ni-NTA Resin Cartridges (ThermoFisher 90098), washed with 20mL of wash buffer (50mM Tris pH 8, 500mM NaCl, 10mM imidazole), followed by elution into 2mL of elution buffer (0mM Tris pH 8, 500mM NaCl, 300mM imidazole). The protein was then dialyzed and concentrated using a size-selection spin concentrator (MilliporeSigma UFC201024).

Protein concentration was determined using a BCA (ThermoFisher 23225) and constructs were analyzed via LC-MS (S14). The liquid chromatography was done using an Agilent 1260 Infinity II system on a PLRP-S column (Agilent PL1912-1500) over an acetonitrile gradient of 5- 95% connected directly to an Agilent 6530 QTOF. Predicted molecular weights were determined using ExPasy ProtParam without the N-terminal Methionine.

#### helixCAM Cell Patterning with Coiled-Coil Coated Plates

First, K562 helixCAM cells were pre-induced for 48 hours with 500ng/mL doxycycline. For testing CC coating concentrations, Ni-coated 96-well plate wells were coated with 100μL of either his-tagged CCs or his-tagged CC-GFP fusions at a range of protein concentrations then incubated for 30 minutes. The CC protein solution was then aspirated and washed with PBS+20mM imidazole (ThermoFisher 88229), then blocked with PBS+20mM imidazole for 30 minutes. The blocking solution was then aspirated, and the pre-induced HelixCAM K562 cells were added in culture media+20mM imidazole+500ng/mL doxycycline for 2 hours. We then washed the plate with culture media twice, followed by imaging and image segmentation to count the number of bound cells. For both dot-based and letter-based dual-CC patterning, the two CC-GFP-His protein solutions used were patterned using a 10uL pipette tip, carefully avoiding contact with the other solution or with the well edge. Subsequent steps are identical to the single CC patterning described above.

#### Imaging

Below is a table summarizing the scopes and setups used for imaging in this work:

| Scope Body | Objectives | Excitation | Ex/Em Setup | Confocal Setup |
| --- | --- | --- | --- | --- |
| Zeiss Axios<br>Observer Z1 | 5X<br>20X | Colibri LED (full<br>spectrum) | DAPI 365/445<br>GFP 470/525<br>YFP 550/605<br>mCherry 560/630<br>Cy5 640/690 | N/A |
| Nikon Eclipse<br>Ti2 | 4X<br>20X | Lumencor SpectraX<br>LED: | DAPI 395/460<br>CFP 440/480 | N/A |
|  | 60X | 395nm<br>440nm<br>470nm<br>510nm<br>550nm<br>575nm<br>640nm | GFP 470/525<br>YFP 510/535<br>mCherry 575/630<br>Cy5 640/700 |  |
| Nikon Eclipse<br>Ti2 | 20X<br>40X | Laser Lines:<br>405nm<br>408nm<br>561nm<br>640nm | DAPI 405/455<br>CFP 405/480<br>GFP 488/525<br>mCh 561/605<br>Cy5 640/705 | W1 Yokogawa<br>Spinning Disk<br>(50µm pinholes) |

Widefield fluorescent microscopy of *E. coli* helixCAM aggregates was conducted using a Nikon Eclipse Ti2 microscope at 60X magnification. Images were taken in the GFP and mCherry channels. Samples were wet-mounted onto a #1.5 coverslip without prior fixation.

Widefield imaging of human cell aggregates (K562, HEK293) was done on either a Zeiss Axios Observer Z1 microscope at 4X or a Nikon Eclipse Ti2 microscope at 20X. For K562 cells, samples were wet-mounted onto a #1.5 coverslip without prior fixation. HEK293 cells were directly grown on a glass-bottom 24-well plate (Mattek P24G-1.5-10-F) and directly imaged. For tiled images composing multiple fields of views, images were taken with a 10% overlap and stitched using the Nikon Elements software.

Widefield imaging of CC-patterned helixCAM K562 cells was done at 4X on a Nikon Eclipse Ti2 microscope. The objective was oriented to the center of the well, and a 2x2 tile was captured and stitched with Nikon Elements, capturing the entire 96-well well surface.

Confocal imaging of K562 cell aggregates and HEK293 cell targeting was done with a Nikon Eclipse Ti2 microscope coupled with a Yokogawa W1 Spinning Disc. In the case of K562 cell aggregates, aggregates were allowed to settle, media was removed, then cells were gently resuspended in melted low melting point agarose (ThermoFisher 16520050) dissolved into culture media and pre-equilibrated to 37°C, then gently dropped onto the glass coverslip of a 24- well glass-bottom plate (Mattek P24G-1.5-10-F). The agarose was allowed to solidify at room temperature for 15 minutes, followed by confocal imaging. 3D reconstruction for 2-color images was done in ImageJ. 3D reconstructions for 3+ color imaging and slice view were created in Nikon Elements.

### QUANTIFICATION AND STATISTICAL ANALYSIS

#### Statistical Summaries and Tests

In Figures 4B, each square represents a mean Δt_50_ value with three replicates. In Figure 4C, the same mean values from Figure 4B are used for the y-axis. R^2^ is the coefficient of determination, and the p-value was determined via an F-statistic test against a constant model. Both values are obtained through MATLAB’s fitlm function. In Figure 4D, the values are the same as in Figure 4B, and error bars are standard deviations. In Figure 4E, values are Δt_50_ values normalized by the max observed Δt_50_ value of that set. The mean of two reads is reported along with standard deviation as error bars. In Figure 7B and Figure 7C, the values are cell counts normalized by the maximum cell counted in that channel and run. This normalization is used to control for variability in cell viability and exposure differences that may confound the absolute count, which is reported in S15 and does not change the interpretation of the findings. The values are reported as medians over four replicates, due to the tendency of small changes in the washing process of producing outliers, and error bars are standard deviations.

#### Next-Generation Sequencing Data Analysis and CC Candidate Selection

FASTQ files from NGS were inspected for quality and then trimmed using Sickle (https://github.com/najoshi/sickle). Trimmed FASTQ files were read into Python via the SeqIO function from the BioPython package (https://biopython.org/) [55]. For the *E. coli* screen, CC candidate sequences were identified, extracted from the read, and paired via an in-house script. CC pairs were then assigned a pair score (S7) which evaluates their frequency of on-target interactions against the frequency of off-target interactions and tuned with a weight factor W. Top 30 hits using a W of 0, 0.01, and 0.1, along with the highest frequency CC candidates, were selected for a total of 102 candidate pairs. From the yeast SynAg screen, fused barcodes representing the two interacting CCs were mapped. CC pairs were then evaluated for enrichment (a function of frequency) and orthogonality (the percentage of on-target interactions divided by all observed interactions), and two pairs (four CCs) with the highest enrichment and orthogonality were selected for characterization in human cells.

#### Image segmentation for measuring interaction frequency and patterning

All image processing and analysis in this work were done downstream of CellProfiler [56]–[59] image segmentation and quantification. For stitched 3-pair fluorescent imaging panels, cells are segmented using raw 12-bit tiff files from each channel using the IdentifyPrimaryObjects module. Exact settings were tuned manually, but primarily using the Otsu 3-Class classification (with the middle class as foreground). For stitched cell patterning images, a circular mask is first applied to remove the signal from the walls of the well.

Illumination correction (CorrectIlluminationCalculate and CorrectIlluminationApply) was applied to reduce the difference in the absolute background due to stitching of 4X images, and IdentifyPrimaryObjects was subsequently used to count the number of cells. Despite best efforts, cells in the overlap region of the stitching were often missed for channels with lower signal intensity, leading to our presentation of the data as normalized cell counts.

#### Image-based Interaction Frequency Analysis Workflow

A detailed explanation of this workflow can be found in S5. In short, cells were segmented and quantified using CellProfiler then imported into MATLAB. The cells in individual channels are merged into a consensus list of cells based on an empirically selected distance threshold. The mean fluorescence of each cell in the list across the five measured channels was calculated, then used as parameters for supervised classification of the cells into the six cell types. Another distance threshold was empirically selected by comparing centroid distances against cell diameters, and this threshold was applied to determine cells that are interacting. Interactions between pairs of cell types are then tallied, then normalized by the sum of the number of interactions in each row and column to obtain the frequency table.

#### HCSRA Processing Workflow

A detailed explanation of this workflow can be found in S11. In short, kinetic OD650 read data over 15 minutes is first imported into MATLAB. For each well, the OD650 is normalized to between 0 to 1, which removes the variance from changes in growth rate affecting the raw OD650 value. The time it took for each sample to reach 0.5 of the normalized OD650 was recorded, which we termed t_50_. For each experiment, both an uninduced and an induced well containing identical cell types and quantity were seeded, and the t_50_ measured for the induced well was subtracted from that of the uninduced well to arrive at a Δt_50_, a measurement of the magnitude of the effect of inducing helixCAM expression in the cells that positively corresponds with increased affinity.

