## Supplemental Figures for "helixCAM: A Platform for Programmable Cellular Assembly in Bacteria and Human Cells"

Supplemental Table of Content:


- S1 – Detailed construct design and plasmid map of bacterial, yeast, and human helixCAM
- S2 – Uncropped images of *E. coli* Z17/Z18, K562 Z17/Z18, P3/AP4, and P9/AP10
- S3 – Sedimentation rate for original three helixCAM pairs (Z17/Z18, P3/AP4, P9/AP10)
- S4 – Full stitched image and channel images of Z17/Z18/P3/AP4/P5/AP6 co-culture
- S5 – Processing Pipeline for Imaging-based cell interaction quantification
- S6 – FACS distributions of GFP intensity across paired sGFP libraries
- S7 – Pair score formula for *E. coli* CC screen
- S8 – Design of yeast helixCAM pre- and post- SUMO addition
- S9 – Large sg30$^{mCherry}$+sg61$^{eGFP}$ aggregate image with scale bar
- S10 – helixCAM-induced aggregation visible without magnification
- S11 – Analysis workflow for HCSRA
- S12 – Full stitched image and channel images of P9/AP10/sg30/sg61/sg83/sg88 co-culture
- S13 – helixCAM-induced K562 binding to HEK293 cells
- S14 – Design of CC-His, CC-GFP-His, gel, and mass spectrometry
- S15 – Absolute Counts for CC-His and CC-GFP-His patterning
- S16 – Dot-shaped patterning of K562 helixCAM cells using CC-GFP-His
- S17 – Table of Key Protein Sequences

S1 – Detailed construct design and plasmid map of bacterial, yeast, and human helixCAM

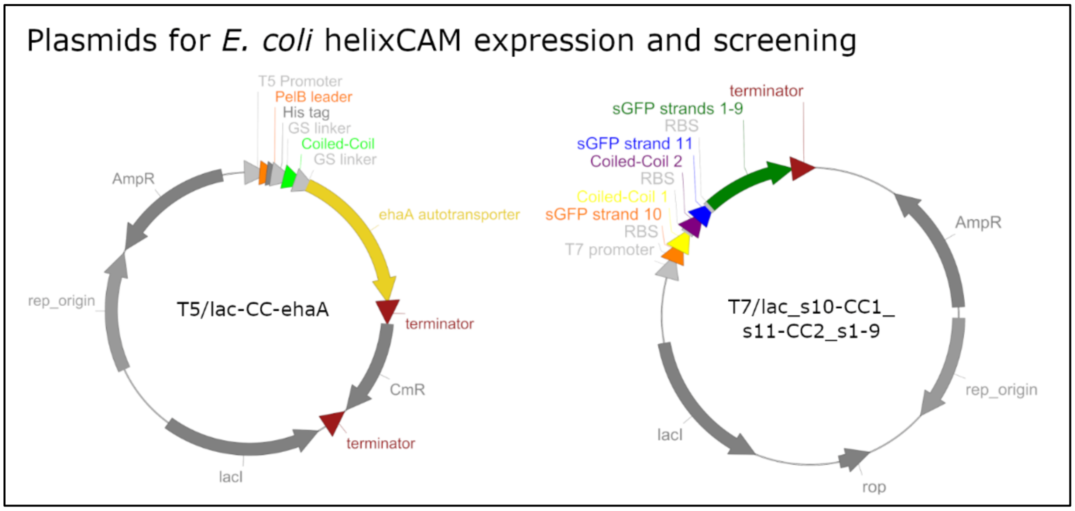


Left: Generic plasmid design for *E. coli* helixCAM expression. The helixCAM consists of a PelB leader sequence, followed by a His-tag, the coiled-coil flanked with glycine-serine linkers, and the ehaA autotransporter protein. The *E. coli* helixCAM expression is controlled by a T5/lac promoter.

Right: Generic plasmid design for the all-in-one tripartite split-GFP plasmid used for stage 1 of coiled-coil library screening. The three components are linked polycistronically under the T7/lac promoter, each with its own ribosomal binding site. The first component is composed of the first coiled-coil fused to the C-terminus of the split GFP $\beta$-strand 10. The second component is composed of the second coiled-coil fused to the N-terminus of the split GFP $\beta$-strand 11. Finally, the third component is the partial the split GFP $\beta$-barrel consisting of strands 1-9. The three components are built on a single plasmid in order to simplify downstream sequencing characterization of positive hits.

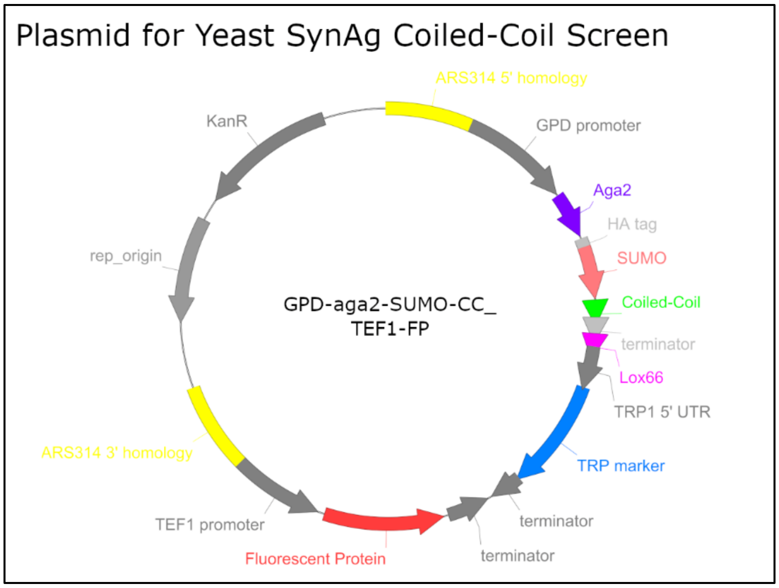

Generic plasmid design for expression of aga2-SUMO-coiled-coil for yeast SynAg assay. The coiled-coil is presented on the cell wall through the fusion to SUMO and aga2 and constitutively expressed under the strong GPD promoter. On the TEF1 promoter, a corresponding fluorescent protein (mCherry or mTurquoise) is expressed. A lox site is present in order to allow Cre to fuse the two distinct plasmid in mated yeast cells. For the library screen, a similar design is used, but rather than a fluorescent protein, either a lysine or a leucine auxotrophic marker was used to allow for complementary auxotrophic selection.

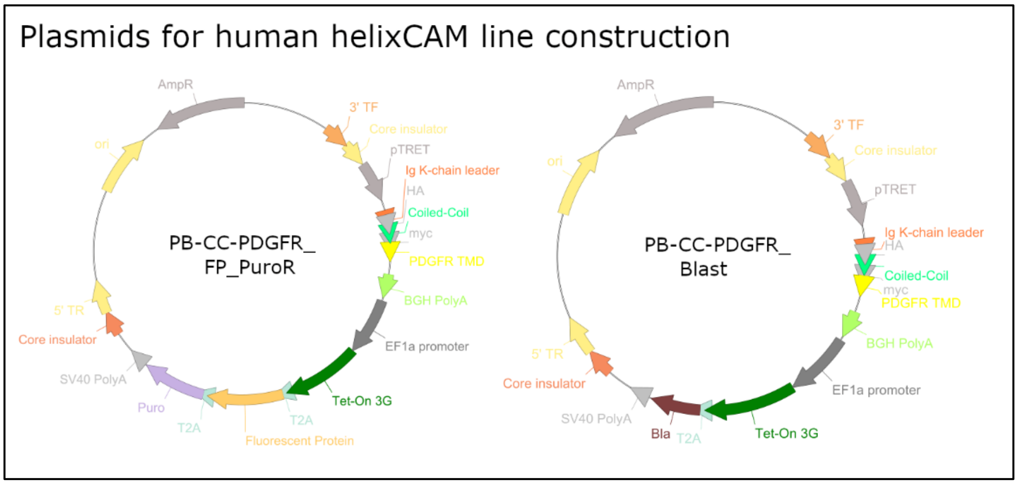


Left: Generic plasmid design used for most human helixCAM lines made in this work. The helixCAM consists of the Ig K-chain leader peptide fused the coiled-coil flanked on either side by an HA tag on the 5' and either a myc or a His tag on the 3', followed by the transmembrane domain of PDGFR. This is driven by the pTRE promoter. The Tet-On activator, fluorescent protein, and puromycin resistance genes are all constitutively expressed polycistronically under a single EF1a promoter.

Right: The plasmid design for blasticidin-selection helixCAM for constructing dual-helixCAM lines. The general design resembles that of the left design, aside from the removal of the fluorescent protein and replacement of the puromycin resistance gene with a blasticidin resistance gene, still polycistronically expressed under EF1a.

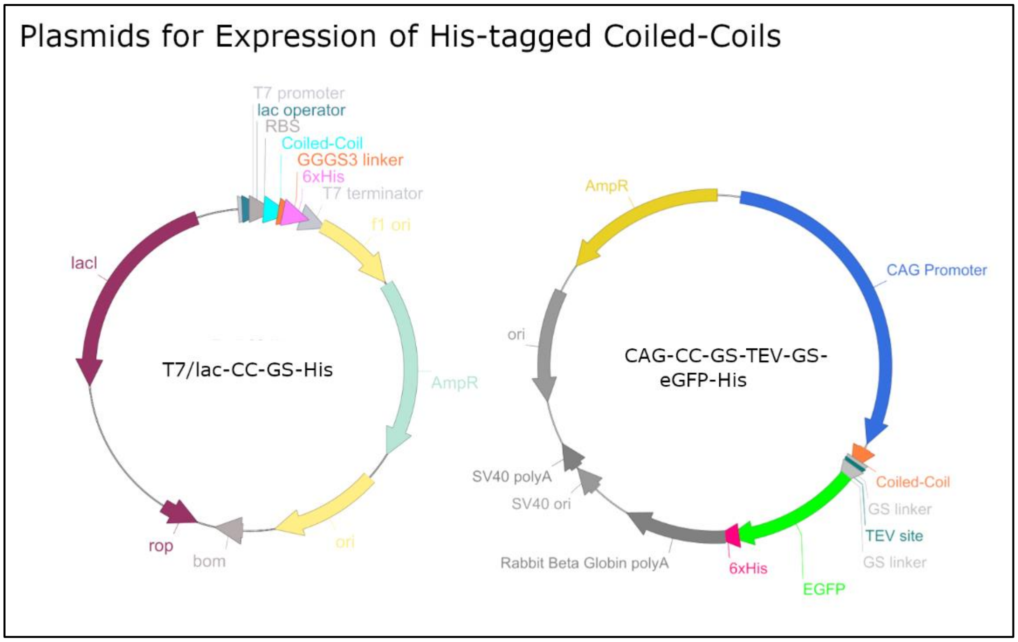


Left: Design for histidine-tagged coiled-coil plasmid for expression in *E coli*. The coiled-coil is tagged with a 12 amino acid glycine-serine linker (3xGGGS) followed by six histidine residues, and expressed under a T7 promoter and tuned with a lac operator.

Right: Design for an eGFP and histidine-tagged coiled-coil plasmid for expression in human cells. The coiled-coil is fused to a Tobacco Etch Virus cleavage site flanked by short glycine-serine linkers (GGSGGG), followed by a full-length eGFP protein, and terminated C-terminally with six histidine residues. The fusion protein is expressed under a CAG promoter for maximal expression.

S2 – Uncropped images of *E. coli* Z17/Z18, K562 Z17/Z18, P3/AP4, and P9/AP10

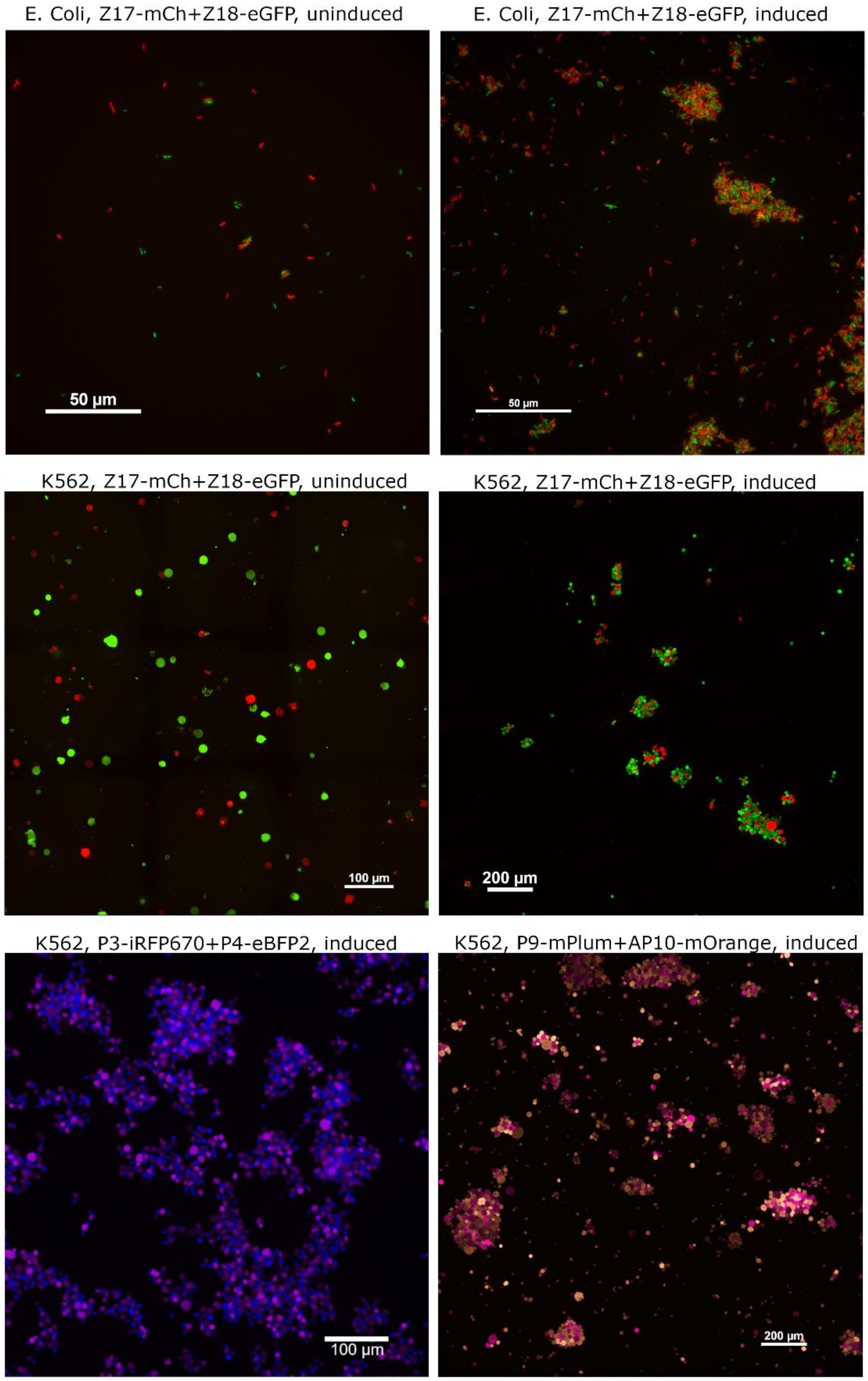


Images presented in Figure 1 and Figure 2 are shown here uncropped to demonstrate representativeness of cropped images.

S3 – Sedimentation rate for original three helixCAM pairs (Z17/Z18, P3/AP4, P9/AP10) in *E coli.*

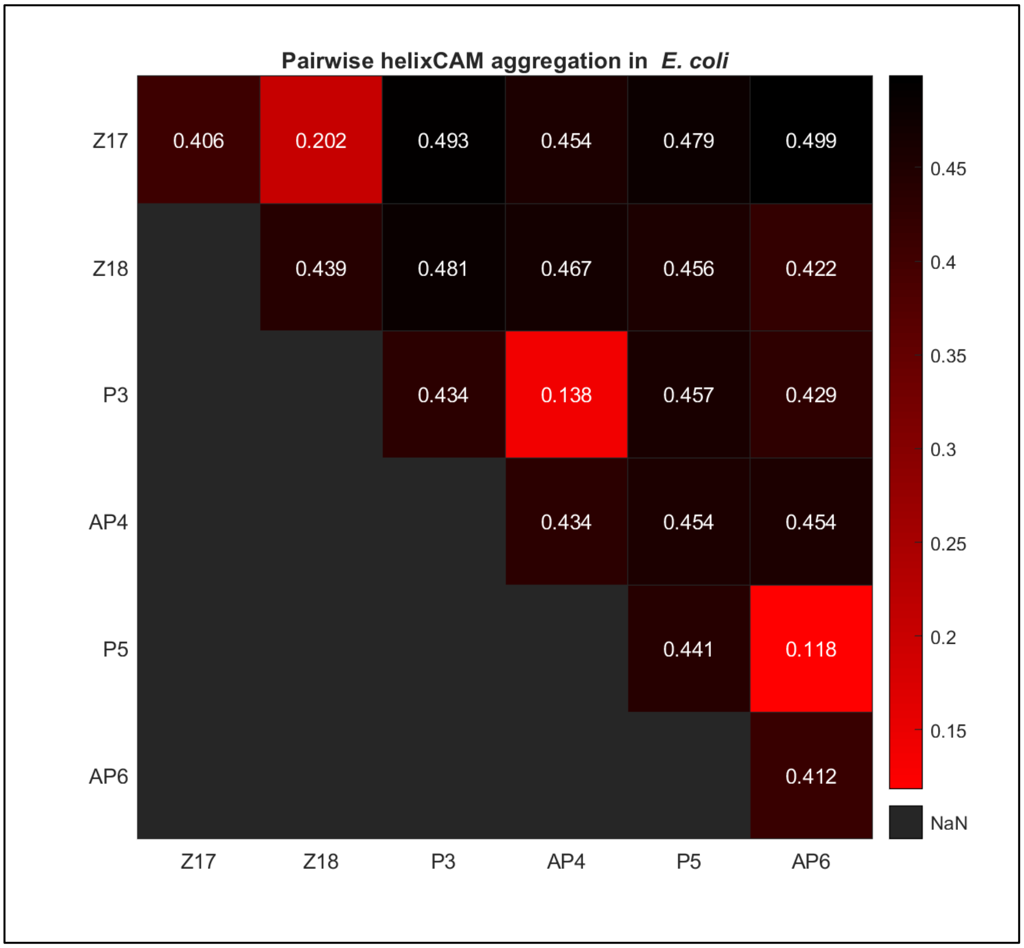


Sedimentation rate of helixCAM-expressing *E. coli* for self- and pairwise interactions. The above heatmap shows the average OD600 across three replicates for each pairwise helixCAM interaction in *E.coli* after four hours of induction and incubation. For the *E. coli* sedimentation assay, the top quarter of the 5mL culture is collected, and the OD600 is immediately measured. Unlike the HCSRA (Figure 4B), lower values indicate faster sedimentation, and thus higher aggregate size and affinity.

S4 – Full image and split channels used for imaging-based interaction quantification of Z17/Z18/P3/AP4/P9/AP10 co-culture

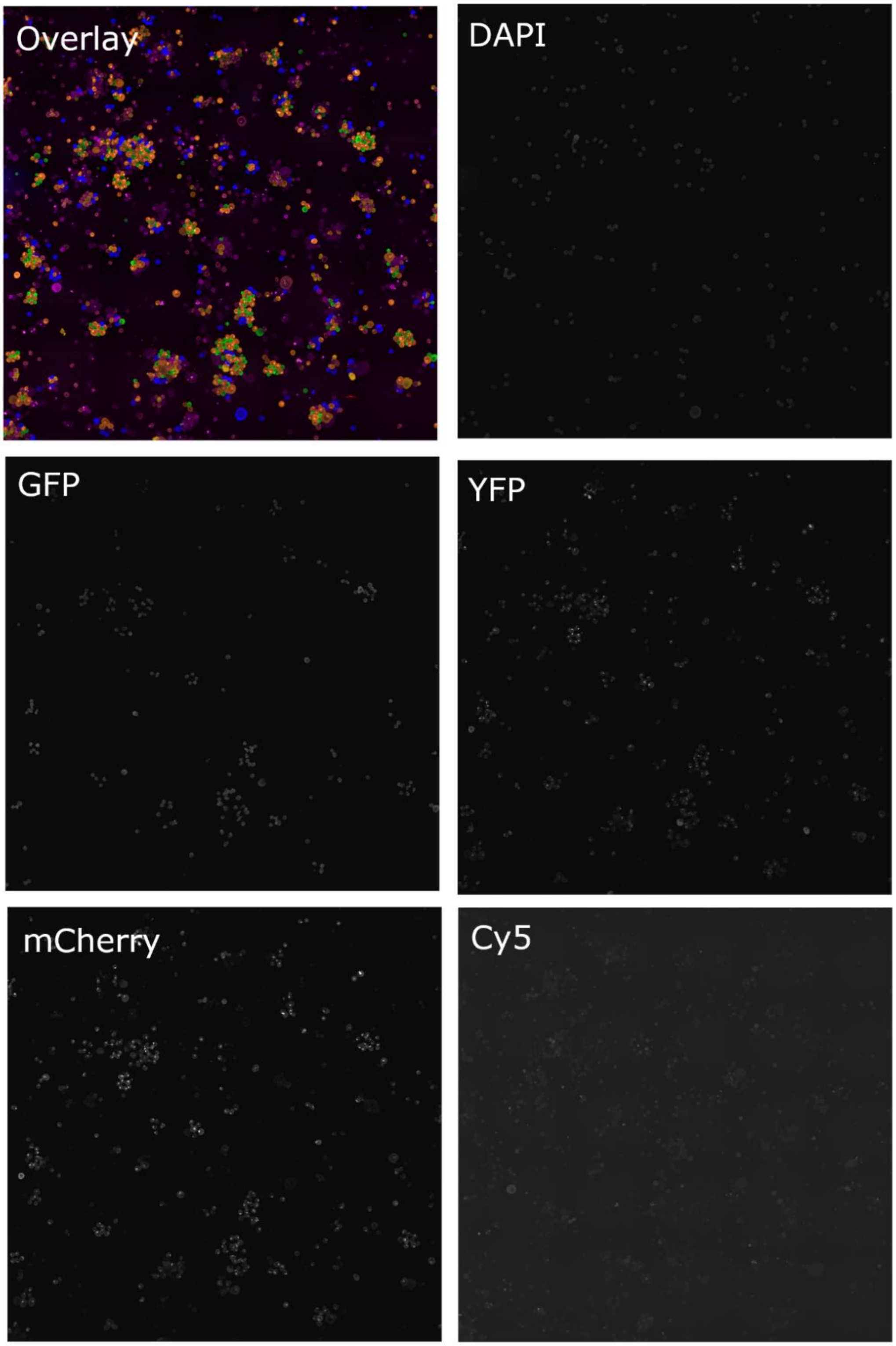

The full stitched image used to calculate the imaging-based interaction frequency (Figure 2D) is shown here. The image was taken using a 6x6 tiling at 20X using a Zeiss Axio Observer Z1 across the five channels shown (DAPI, GFP, YFP, mCherry, and Cy5) with a 10% overlap, followed by stitching in Nikon Elements.

S5 – Processing Pipeline for Imaging-based cell interaction quantification

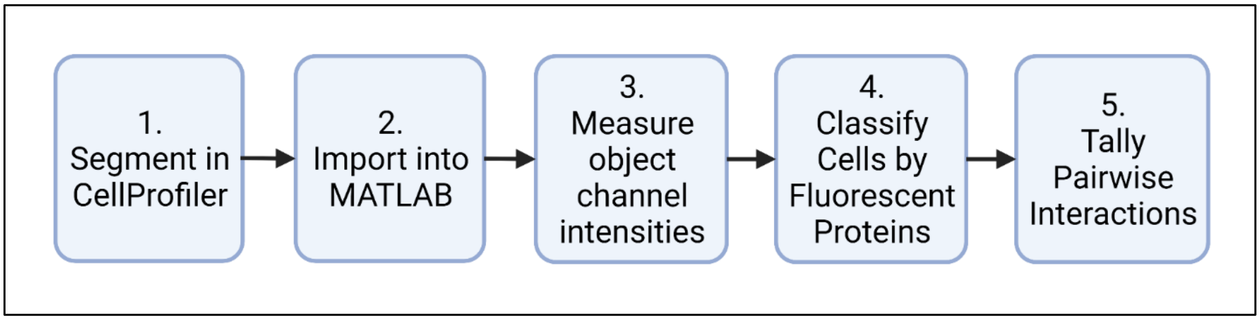


Our image-segmentation based method for characterizing cell-cell interaction frequency consists of five general components. First, cells are segmented using CellProfiler. Next, the object data is imported into MATLAB. At this point, we import the raw image data from each channel, and measure average intensities for each object in each channel. This is followed by a supervised classification of the objects into the six fluorescent protein-expressing cell types. Finally, interactions are assigned based on distance and assigned to pairs.

1. Image segmentation and Object Identification in CellProfiler

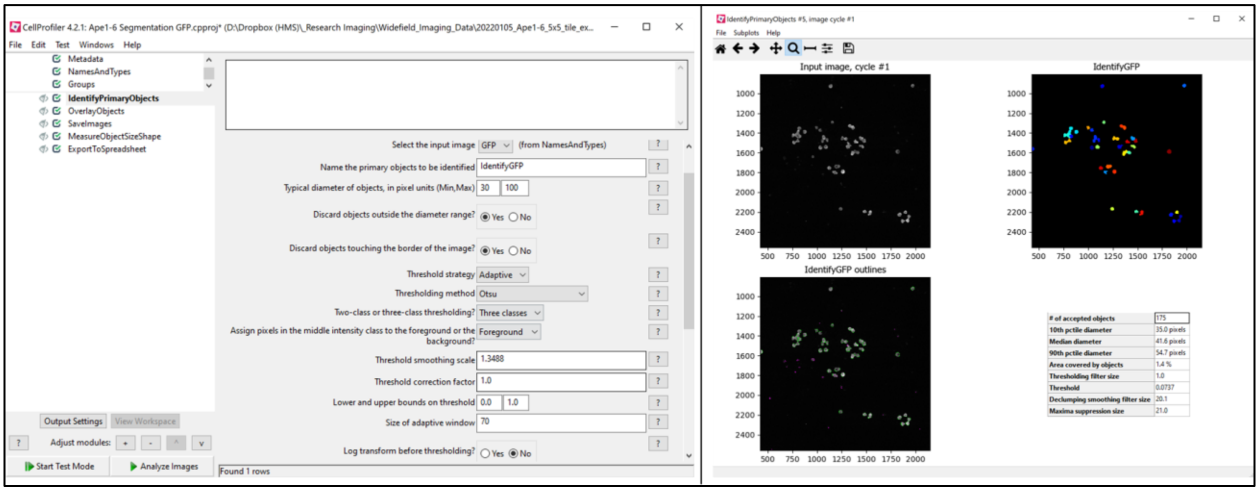


The stitched .tif image in each of the five channels (DAPI, GFP, YFP, mCherry, Cy5) is first segmented through CellProfiler, primarily through the IdentifyPrimaryObject module. The exact values change from image to image, but generally a diameter of 30-100 along with adaptive thresholding using Otsu 3-class (middle intensity class assigned to foreground), worked well for most channels (Left). On the right panel, it is possible to see an example of the segmentation obtained for the GFP channel (image is zoomed in for clarity).

2. Object import into MATLAB

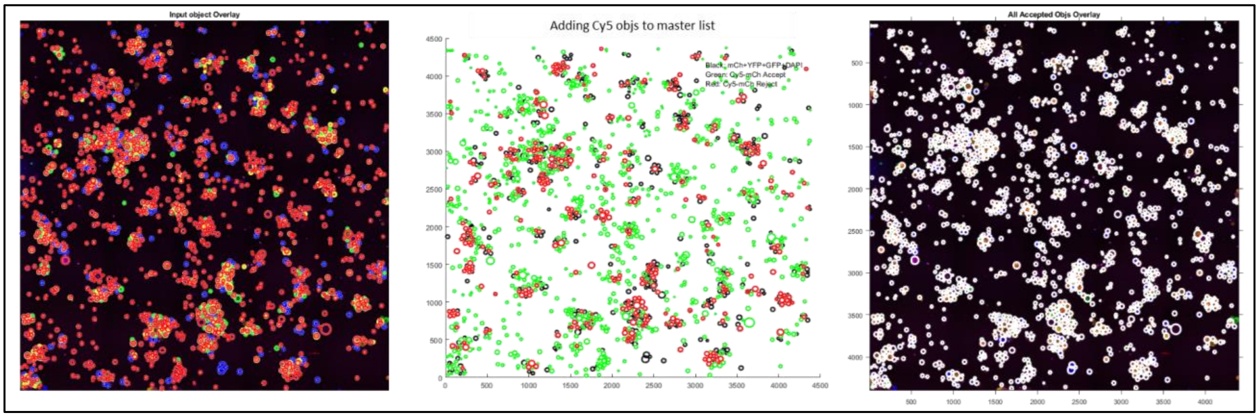


Objects from each channel are first imported into MATLAB. The left pane shows the objects, outlined by channel color, overlaid on the fluorescent image. However, as some fluorescent proteins have signal across multiple channels, it is necessary to identify those cells as a single object during import. To do this, we set a distance threshold based on a histogram of pairwise distances, then sequentially merge the list of objects into a master list from the highest to lowest signal to noise ratio (GFP->DAPI->mCherry->YFP->Cy5). The center pane shows the process of merging Cy5 objects into the list. Black circles show existing master list objects, green circles show newly accepted objects, and red circles show rejected objects. The right pane shows the outline of all master list objects on the fluorescent image.

3. Measure object Intensity

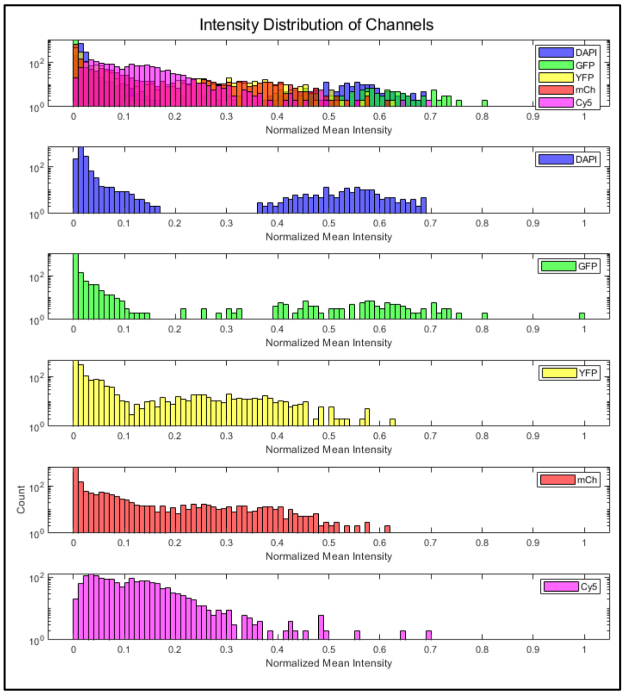


We calculated the fluorescence of each accepted object by determining cell boundaries using their imported X/Y coordinates and diameter, and measuring the mean intensity from the raw channel image data. The histograms on the left shows the distribution of intensities in each channel. It is possible to identify a bimodal distribution for each channel corresponding to cells either positive or negative to that channel.

4. Supervised Classification of objects by fluorescent protein

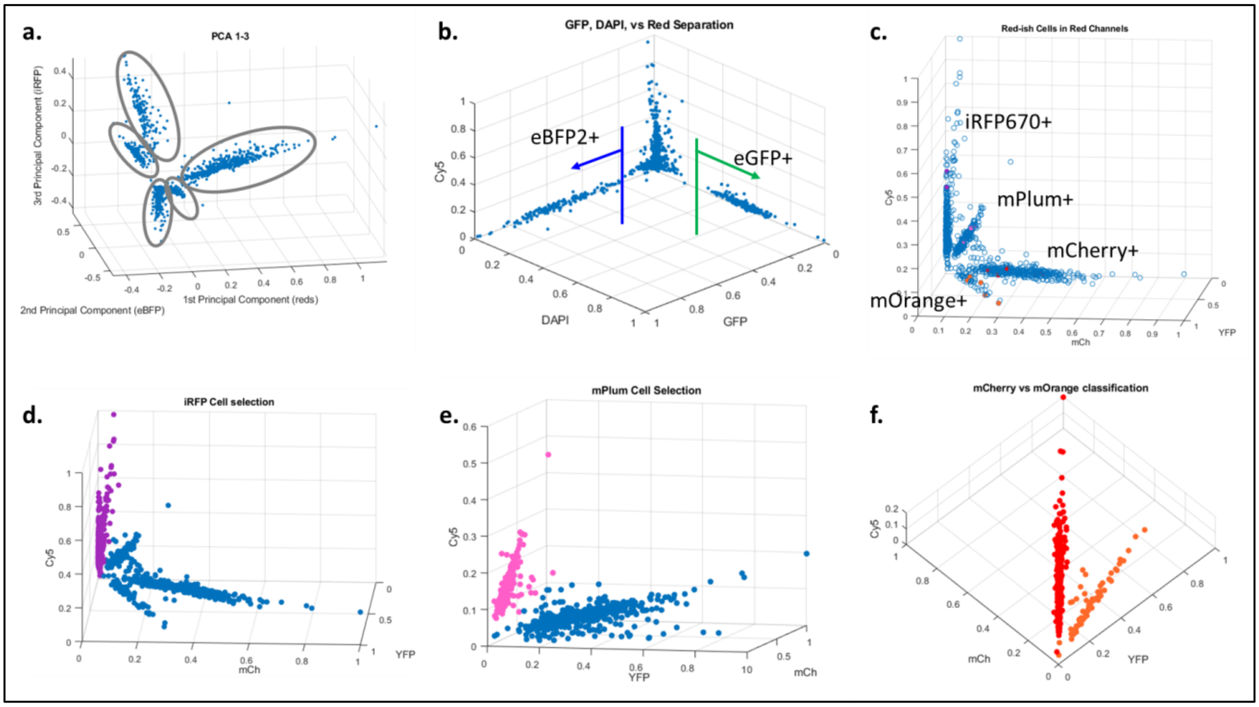


As mentioned, to identify six FP-expressing cell types using five channels, certain cells will be visible in multiple channels. However, through PCA of the mean intensities of the objects, it is possible to see that there are distinct clusters, meaning it is possible to categorize each cell by fluorescence (a). First, we look at the two FPs most distinct in their channels, eBFP2 and eGFP. For these two cell types, a simple intensity threshold can be used for classification. Removing these two populations, we visualize the remaining four cell populations in the YFP, mCherry, and Cy5 channels alongside manually annotated cells to identify which population expresses which FP. We then sequentially classify cells and remove them from the population using an inequality hyperplane defined by two channels (iRFP670: Cy5, mCherry; mPlum: Cy5, YFP; mOrange: mCherry, YFP). The classified cells can be seen in the below graph:

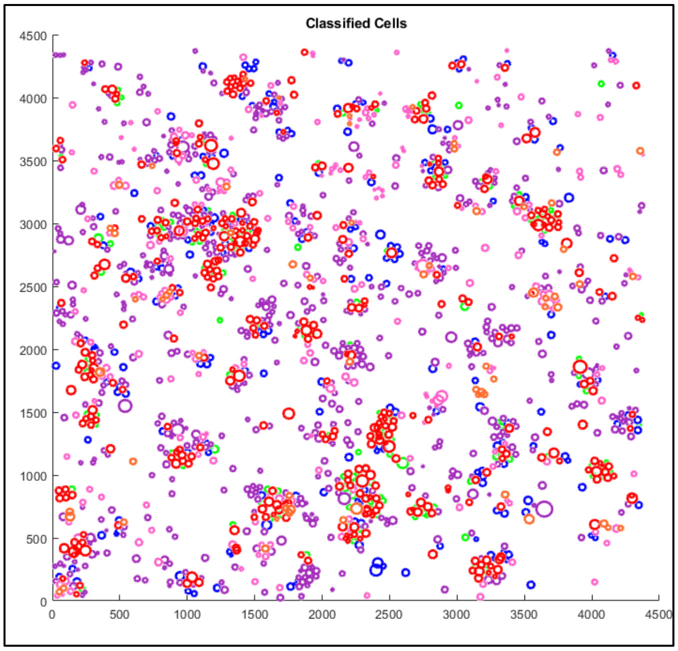


5. Identification cell-cell interaction and tallying of self- and pairwise interactions

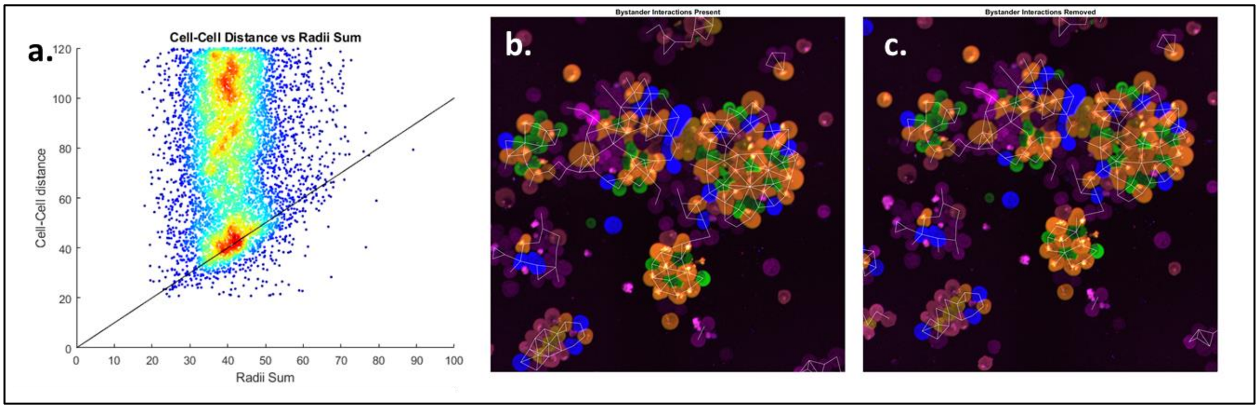


Graphing a heatmap scatterplot of all distances between pairs of cells against the sum of the pairs' radii, it is possible to see a cluster form near the y=x diagonal (a). We define this cluster as "interacting cells," which we can visualize by overlaying on top of the fluorescent image (b). With a purely distanced-based approach, however, the method would also consider cells which happen to be next to each other due to shared binding to a third cell, as a self-interaction. However, this is likely not reflective of the true interactions occurring. To address this, we prune interactions by removing "interactions" between cells that share a third interaction partner. In practice, this filter does not remove many interactions (<10%), and its effects can be seen by comparing (b) and (c).

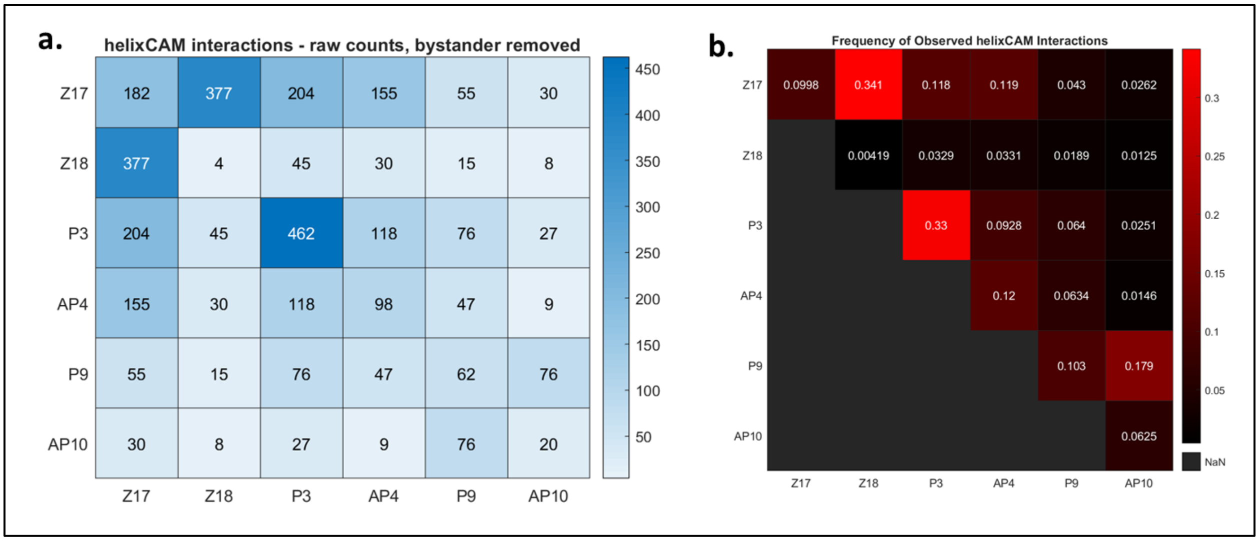


After determining interacting cells, we tally the number of observed interactions for each self and paired interactions as a table (a), then divide each square by the sum of the number of interactions in its row and column to obtain a frequency matrix, which is mirrored across the diagonal (b). We present this data as a black/red heatmap in Figure 2D.

S6 – FACS distributions of GFP intensity across paired sGFP libraries

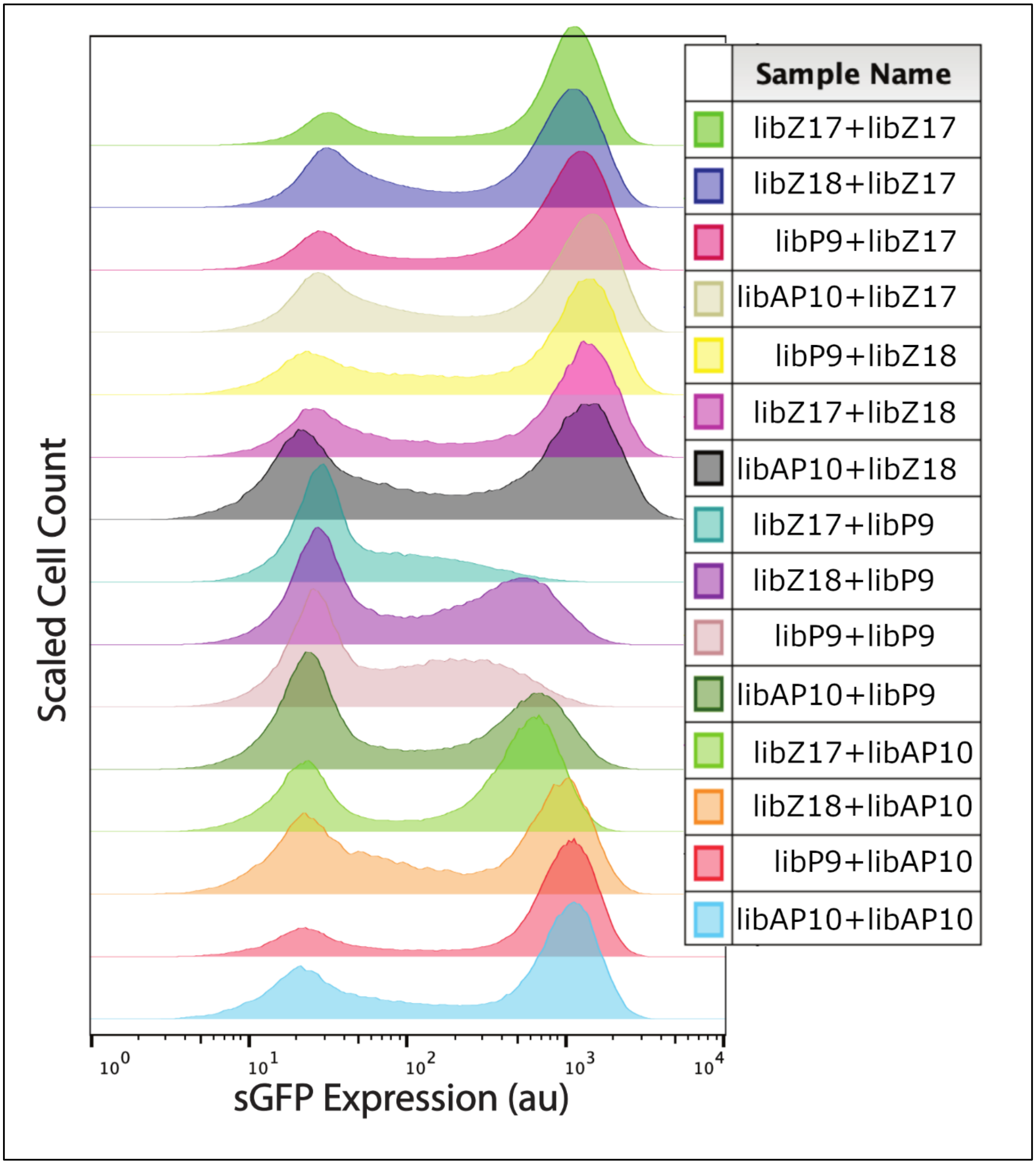


As the higher throughput first stage of our two-stage coiled-coil library screen, we used the tripartite split-GFP approach in *E. coli*. eGFP is split into β-strands 1-9, and β-strands 10 and 11. These three components have low affinity to each other unless β-strands 10 and 11 are brought together by a separate domain, leading to formation of the intact GFP β-barrel and green fluorescence. We built four coiled-coil libraries based on Z17, Z18, P9, and AP10, and fused them to either the C-terminus of strand 10 or N-terminus of strand 11. We then mixed these libraries in pairs, transformed into competent cells, and sorted for GFP-positive cells. All mixed populations demonstrated a bimodal distribution, demonstrating both a functional screen as well as interacting coiled-coil pairs in each pool.

S7 – Pair score formula for E. coli CC screen

$$\text{Pair score} = \frac{f_p}{\underbrace{\left(\sum f_{C_1} + \sum f_{C_2} - f_p\right)}_{\text{Penalizes frequent non-pair binding}} + \underbrace{W(P_1 * P_2 - 2)}_{\text{Penalizes number of unique binding partners}}}$$

$f_p$ = pair frequency
$W$ = weight
$C_1$ = coil one
$C_2$ = coil two
$P_1$ = # of pairs including $C_1$
$P_2$ = # of pairs including $C_2$

A pair score was calculated for each pair of CC candidates to bring into the next stage of selection. The pair score considers two metrics: the frequency of each pair binding with other CCs in the pool, and the number of unique CCs each pair binds. The latter metric is multiplied by a weight factor W, which effectively tunes the score to either favor CCs that bind a smaller set of other CCs strongly, or CCs which bind a large number of CCs weakly. W's used were 0, 0.01, and 0.1, and the top 30 pairs with from each W, along with the highest frequency pairs, were selected for the next round of screening (totaling 102 CC candidates).

S8 –yeast SynAg assay results before and after addition of SUMO

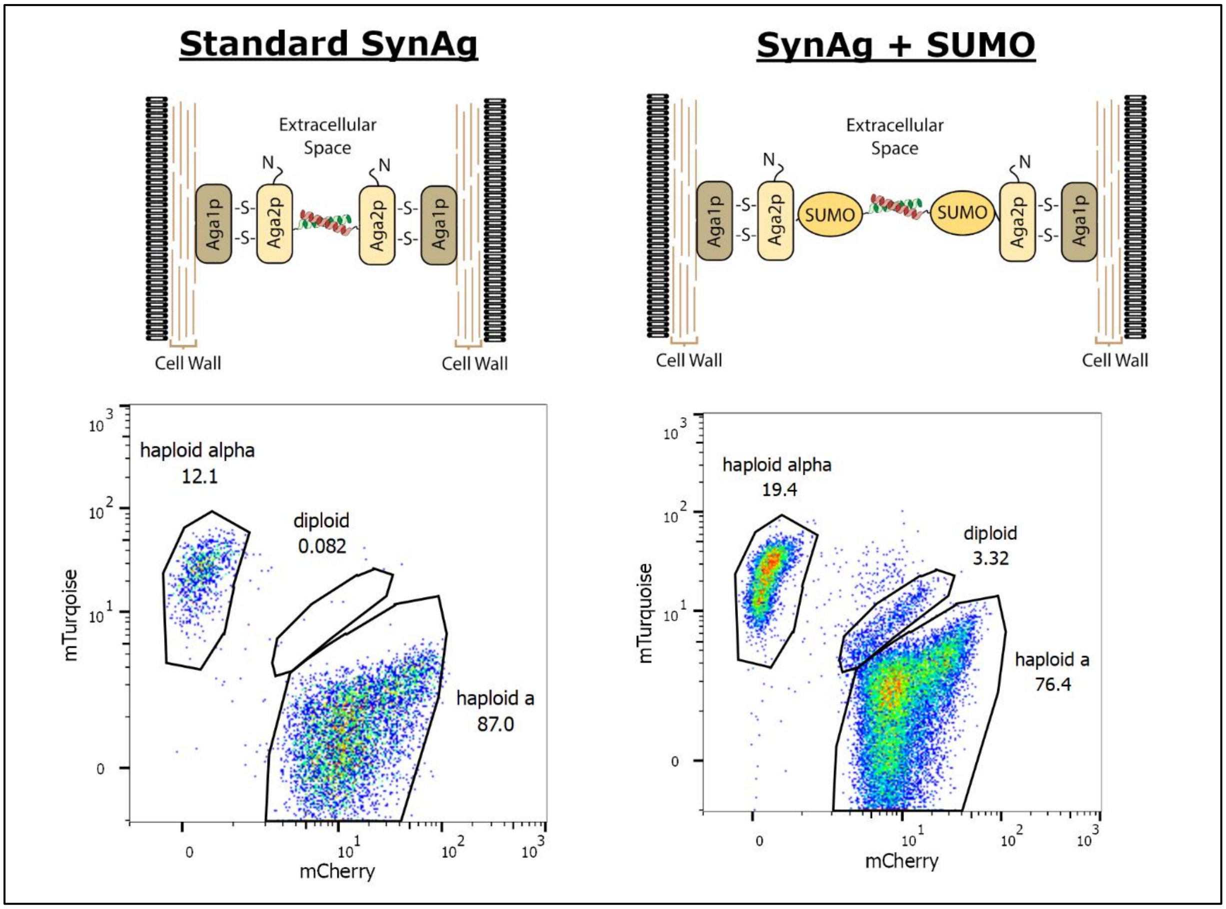


For stage two of our coiled-coil screening approach, we looked to enrich for helixCAM-compatible candidates – coiled-coils amenable to binding when presented on the cell surface. To do this at a high throughput, we used the yeast SynAg assay, which uses surface protein-induced mating of haplotype alpha and haplotype a yeast cells into diploid, leading to cells expressing two markers (either two fluorescent proteins or two auxotrophic markers). We first tested the standard SynAg using Z17 and Z18 with the mCherry and mTurquoise fluorescent proteins. However, we did not observe the expected dual-positive population that successful mating should create. We hypothesized that this was due to trouble with expressing the coiled-coil-Aga2p fusion and localizing it to the yeast cell wall. To resolve this, we tried inserting a SUMO protein domain between the coiled-coil and Aga2p to help improve stability and solubility. This resulted in the formation of a third, dual-fluorescent yeast population, indicating successful yeast mating, so we adopted this design for stage two of our coiled-coil screening process.

S9 – Large sg30$^{mCherry}$+sg61$^{eGFP}$ aggregate image with scale bar

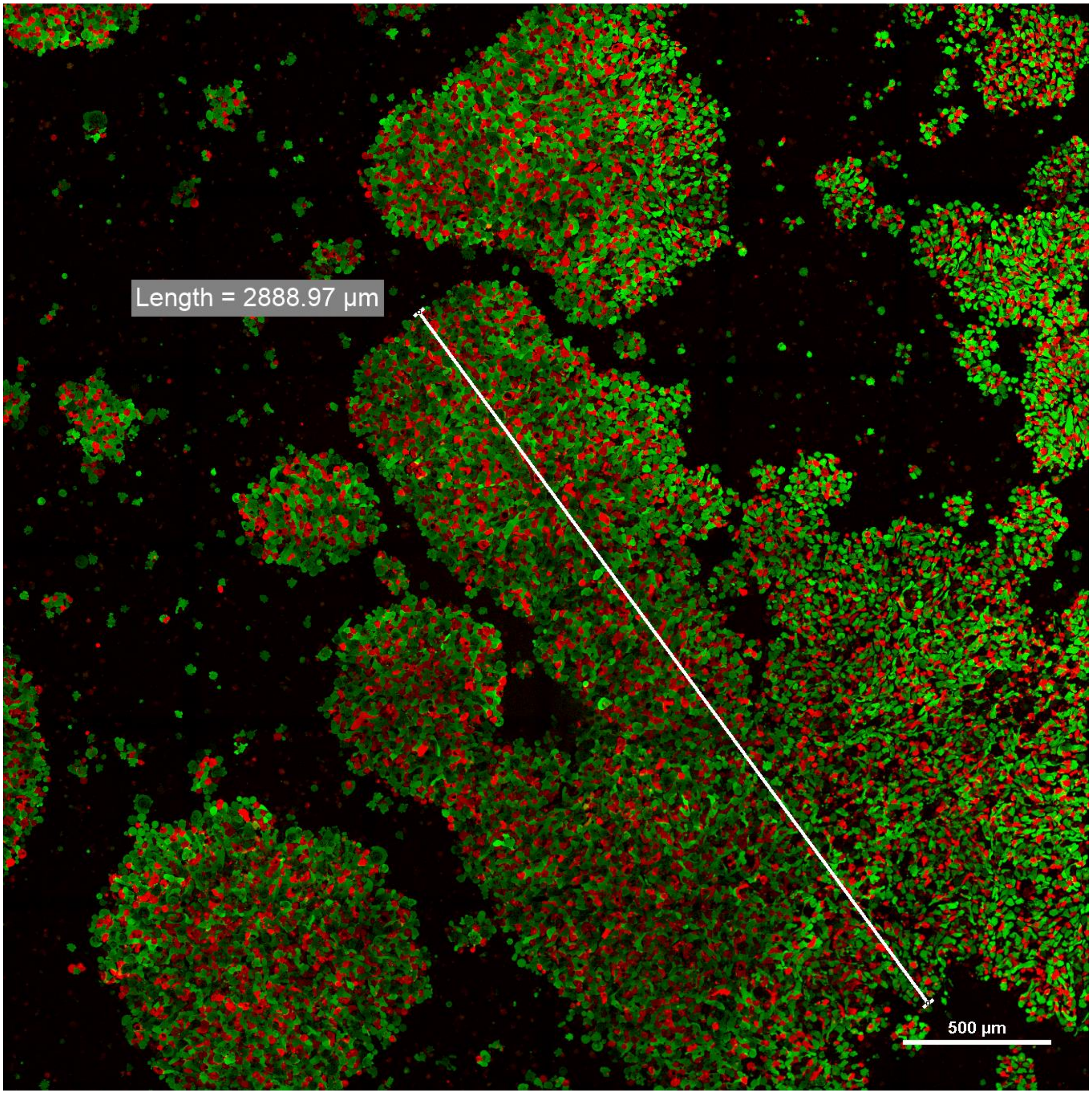


Measurement of large sg30+sg61 aggregation. Sg30 and sg61 K562 helixCAM cells were induced with doxycycline and incubated for 48 hours in a 24-well nonstick plate. At the 48 hour point, large aggregates were visible by eye. 20uL of these aggregates were transferred with a wide-bore pipetted and mounted onto a glass coverslip, followed by widefield imaging at 20X magnification. This image is stitched from 6x6 tiling with 10% overlap, and distance overlay was done in the imaging software (Nikon Elements). When compressed by the coverslip, the largest aggregate size observed was over 2.8 millimeters in length. The ability for helixCAMs to form large aggregates indicates strong potential for use in tissue engineering.

S10 – helixCAM-induced aggregation visible without magnification

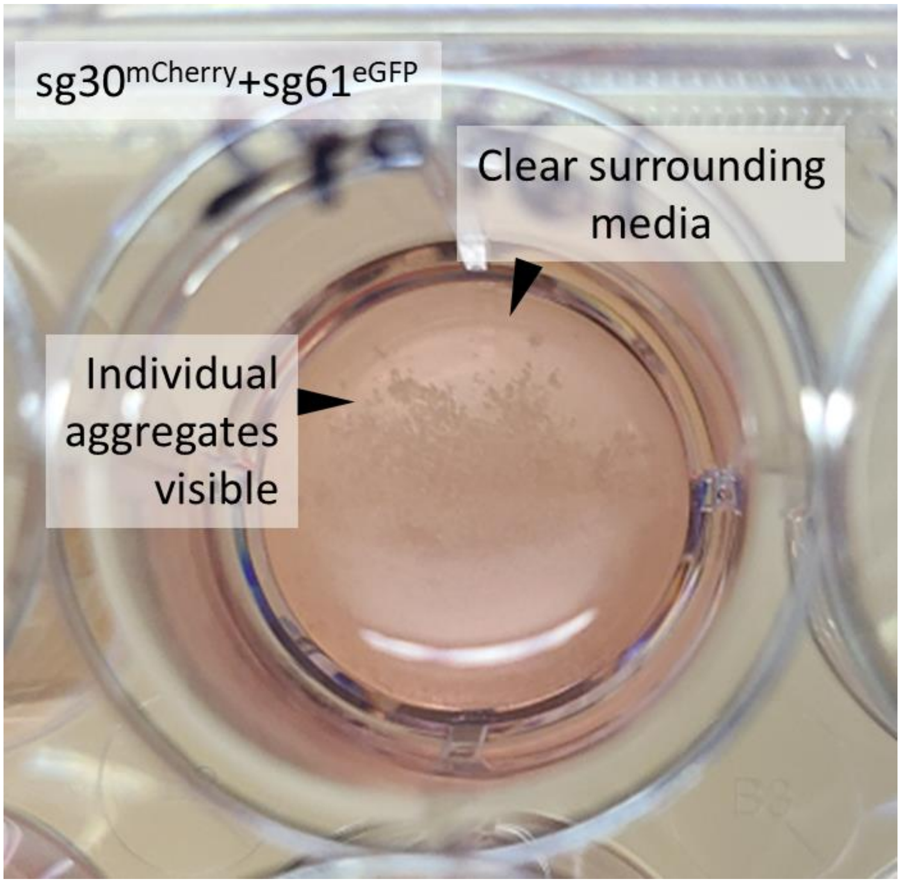


Image of 12-well plate well containing a mixture of sg30$^{mCherry}$ and sg61$^{eGFP}$ cells after 48 hours of induction. Cell aggregates can seen by eye, and image was captured with a cell phone camera at 1X magnification.

S11 – Analysis Workflow for HCSRA

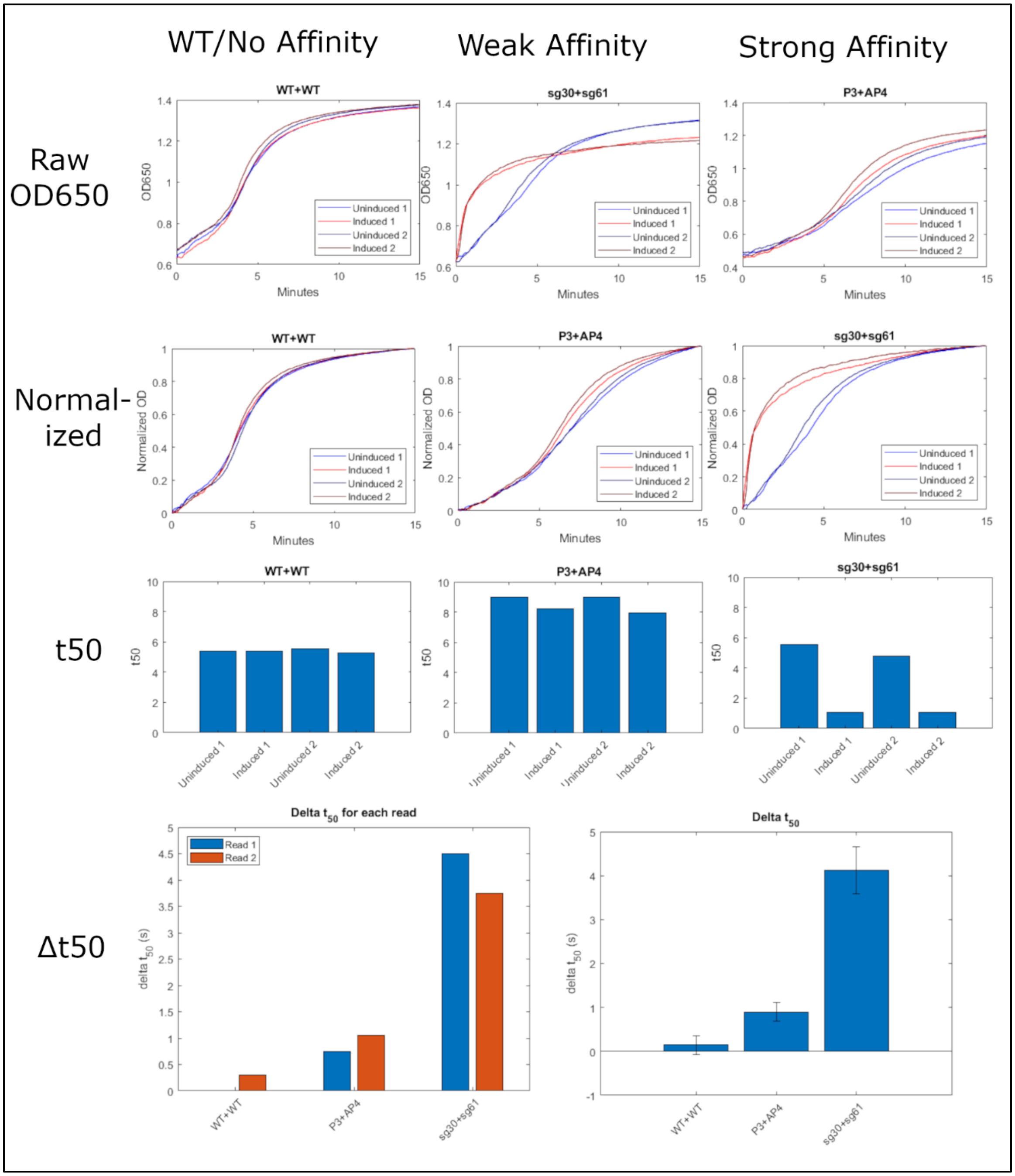


Measurement and analysis workflow for Human Cell Sedimentation Rate Assay. K562 helixCAM cells are mixed together at 1.875e5 cells per population for paired interactions or 3.75e5 for self-interaction in 150uL of DMEM+10% FBS+1% P/S+1ug/mL doxycycline in ultra-low-adhesion v-bottom 96-well plates. The plate is incubated at 37 °C and 5% $CO_2$ with shaking for 48 hours. Plates are then shaken at 900rpm for 15 seconds, immediately followed by OD650 measurement for 15 minutes at 37 °C. The plate is then shaken again and the OD650 is read again, resulting in two reads. The raw data can be seen in the top row. Due to some variability in growth rate between induction and cell types, the OD650 values are normalized by subtracting by the minimum, then divided by the maximum to span a range of 0 to 1 (row 2). From this data, the time point at which the OD surpasses 0.5 is taken, which we refer to as $t_{50}$ (row 3). The $t_{50}$ for the induced condition is then subtracted from the $t_{50}$ from the uninduced condition, leading to a positive value if doxycycline induction led to faster sedimentation and thus larger aggregates, and a negative value if doxycycline induction led to slower sedimentation. Finally, the values are averaged to obtain a final representative $t_{50}$ value for each pairwise and self interaction. As seen in Figure 4c, the values from the HCSRA correlate to interaction frequencies observed through microscopy, indicating that the HCSRA is a simple and effective method for measuring cell-cell affinity.

S12 - Full image and split channels used for imaging-based interaction quantification of Z17/Z18/P3/AP4/P9/AP10 co-culture

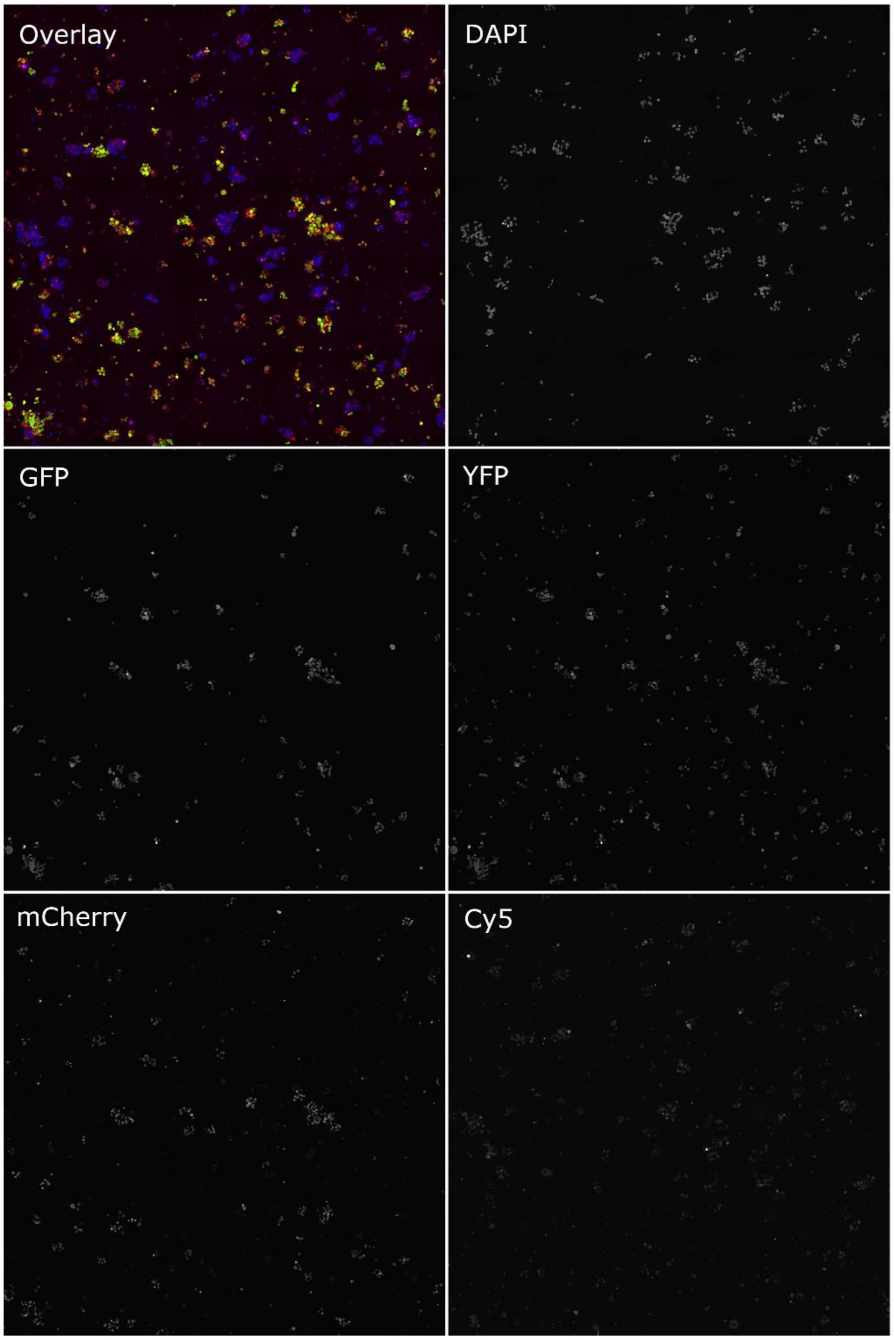


The full stitched image used to calculate the imaging-based interaction frequency (Figure 2d) is shown here. The image was taken using 5x5 tiling at 20X using a Nikon Eclipse Ti2 microscope across the five channels shown (DAPI, GFP, YFP, mCherry, and Cy5) with a 10% overlap, followed by stitching in Nikon Elements.

S13 – helixCAM-induced K562 binding to HEK293 cells

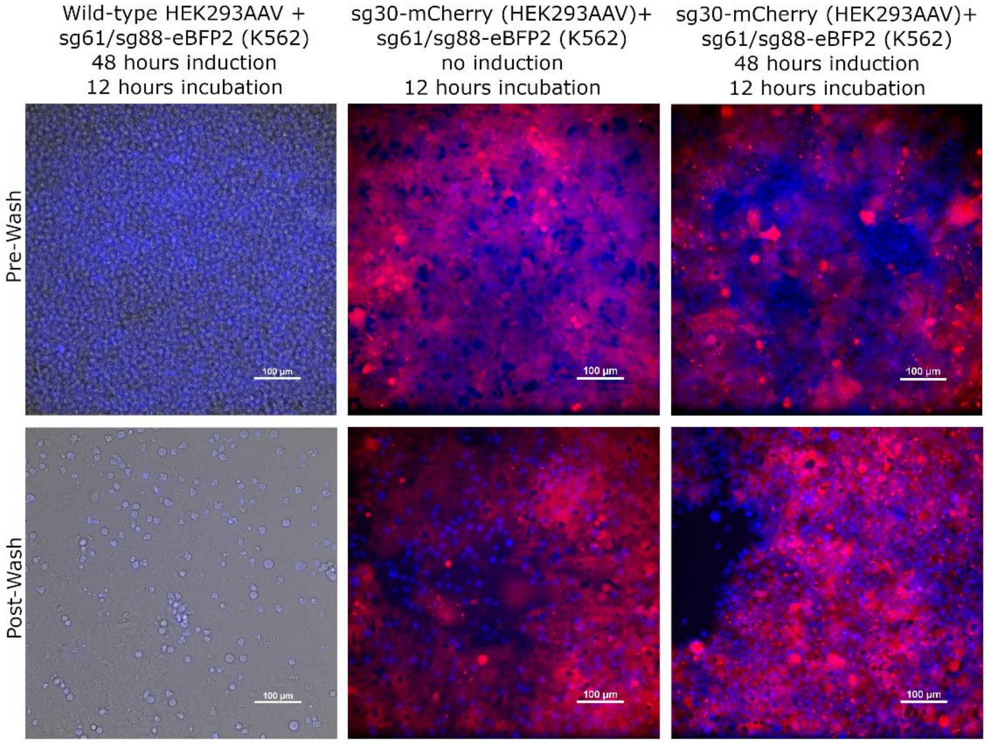


In order to demonstrate the capability of helixCAMs to selectively target natively suspension cells to adherent cells for applications such as immuno-oncology, we tested our sg61/sg88-eBFP2 K562 cells against either wild-type HEK293 cells and sg30-mCherry HEK293 cells, both uninduced and induced. K562 cells are a good fit for this experiment, stemming from lymphoblast lineage, and HEK293 an interesting target – stemming from an embryonic adrenal lineage. The two cell types for each experiment are grown separately to 80% confluency, followed by either no induction (center column) or 48 hours of induction (right column). The suspension cells are then added to the adherent cells, followed by an additional 12 hours of incubation. The cells are imaged prior to wash, then washed twice with culture medium with gentle linear shaking. The cells are then imaged again. Brightfield is only shown for wild-type HEK293 conditions to demonstrate complete coverage of the field of view. While there are some remnant cells in both the wild-type and uninduced sg30-mCherry cell conditions, the induced condition has a significantly denser coat of eBFP2-expressing K562 cells covering the surface of the mCherry-expressing HEK293 cells, demonstrating the ability of helixCAMs to drastically increase binding of suspension cells to adherent cells.

S14 – Design of His-tagged coiled-coils, gel image, and mass spectrometry

<table>
<tr><th>Name</th><th>MW (Da)</th><th></th></tr>
<tr><td rowspan="2">sg30-His</td><td rowspan="2">6118</td><td>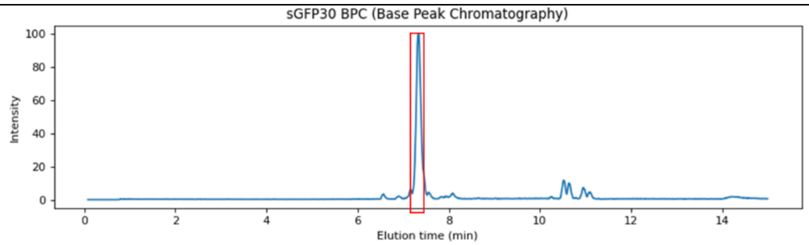
</td></tr>
<tr><td>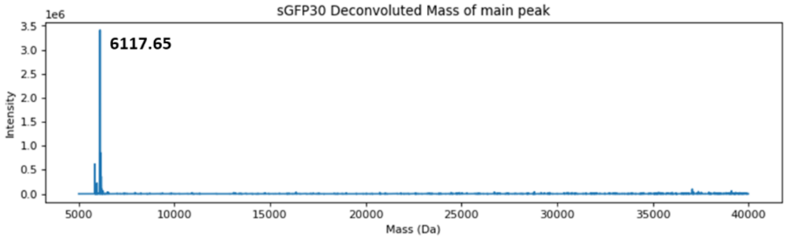
</td></tr>
<tr><td rowspan="2">sg61-His</td><td rowspan="2">6115</td><td>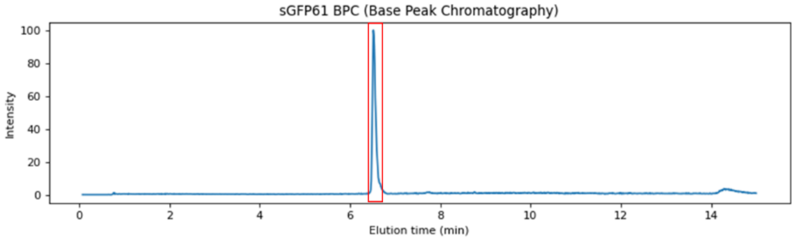
</td></tr>
<tr><td>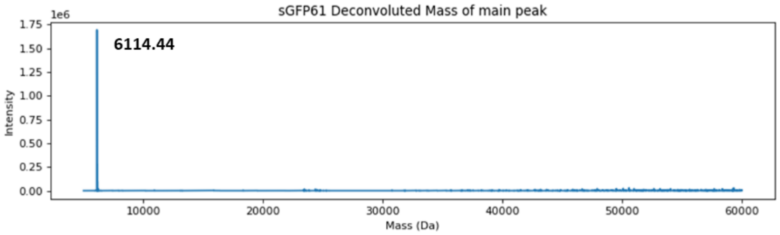
</td></tr>
<tr><td rowspan="2">sg83-His</td><td rowspan="2">6220</td><td>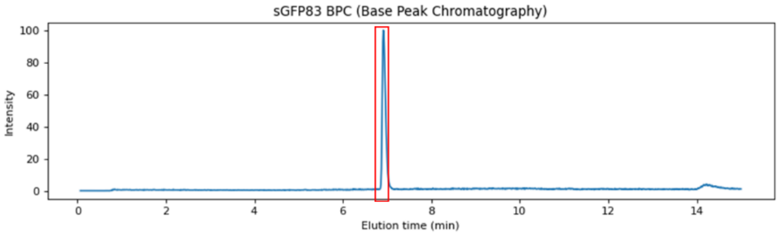
</td></tr>
<tr><td>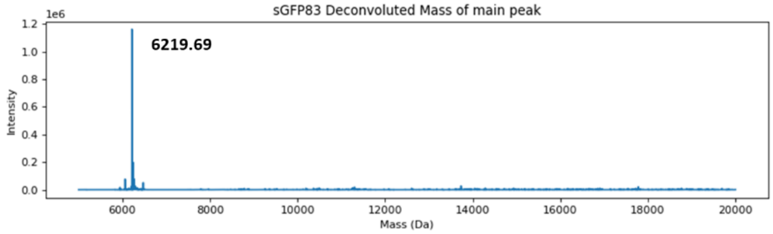
</td></tr>
</table>

<table>
<tr><td rowspan="2">sg88-His</td><td rowspan="2">6188</td><td>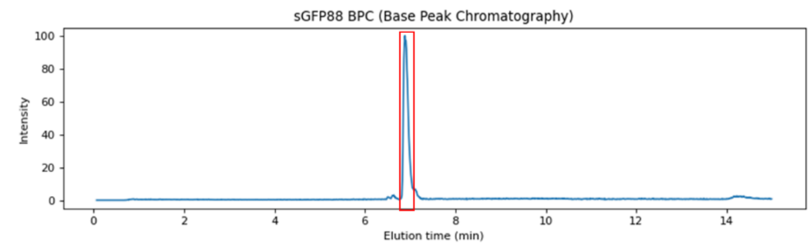
</td></tr>
<tr><td>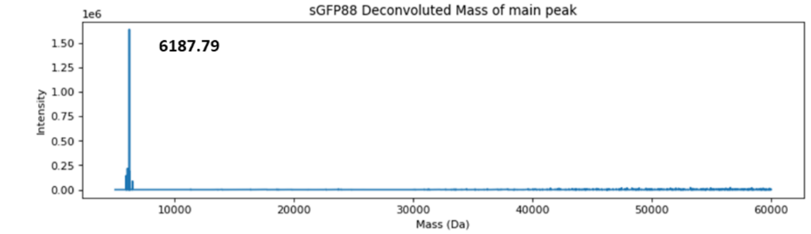
</td></tr>
<tr><td rowspan="2">sg30-GFP-His</td><td rowspan="2">33853</td><td>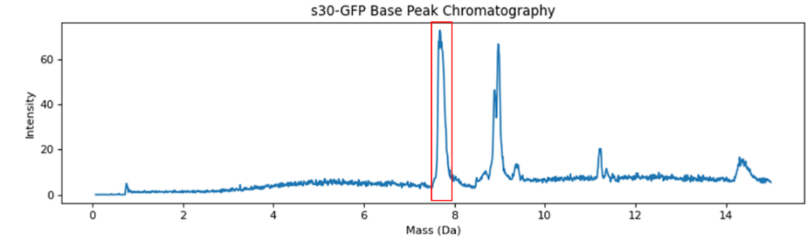
</td></tr>
<tr><td>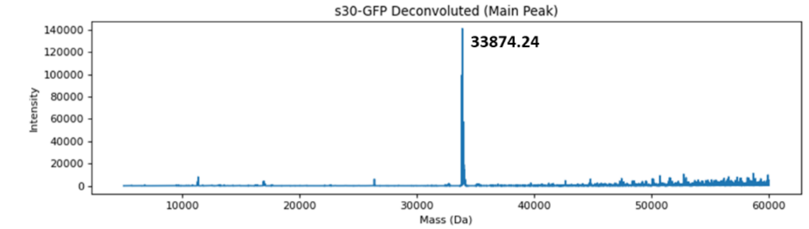
</td></tr>
<tr><td rowspan="2">sg61-GFP-His</td><td rowspan="2">33850</td><td>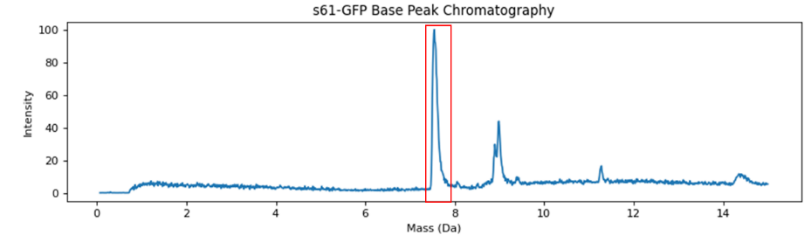
</td></tr>
<tr><td>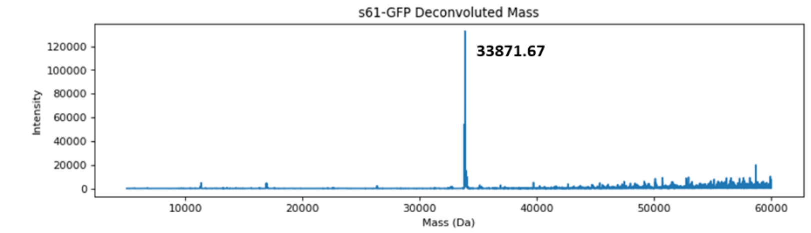
</td></tr>
</table>

<table>
<tr><td rowspan="2">sg83-GFP-His</td><td rowspan="2">33955</td><td>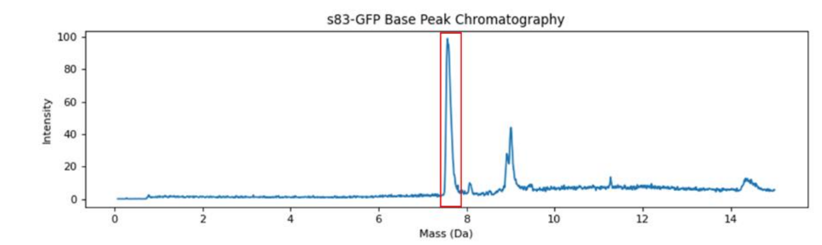
</td></tr>
<tr><td>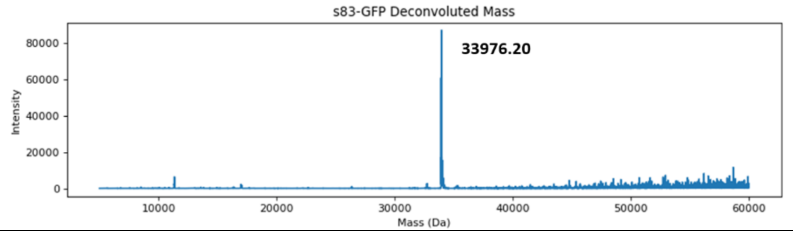
</td></tr>
<tr><td rowspan="2">sg88-GFP-His</td><td rowspan="2">33923</td><td>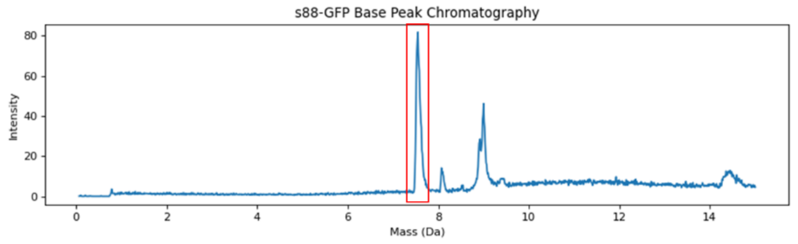
</td></tr>
<tr><td>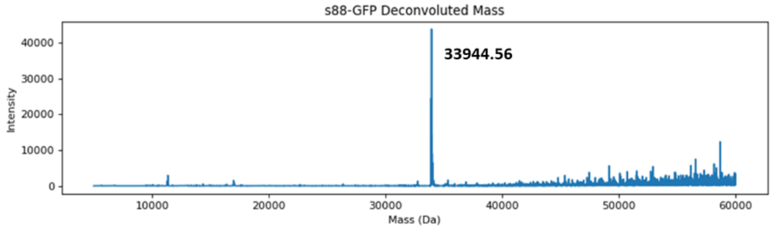
</td></tr>
</table>

Coiled-coils tagged with either 6 histidine residues or both GFP and 6 histidine residues were expressed using *E. coli* or HEK293 cells respectively and subsequently analyzed by liquid chromatography (top trace) and mass spectrometry (bottom trace). The CC-His proteins (first four) were reasonably pure, with only one major peak in the LC trace and likewise in the MS trace corresponding to the expected molecular weight. The CC-GFP-His proteins had a side peak that eluted later in LC, which, when run through MS, shows the same molecular weight, leading us to conclude that it represents the unfolded CC-GFP-His protein. The resulting MS measurement of molecular weight for CC-GFP-His proteins were all higher than expected by 22 Da, which we believe to be an addition of a sodium ion from the PBS buffer used for purification.

S15 – Absolute Counts for CC-His and CC-GFP-His patterning

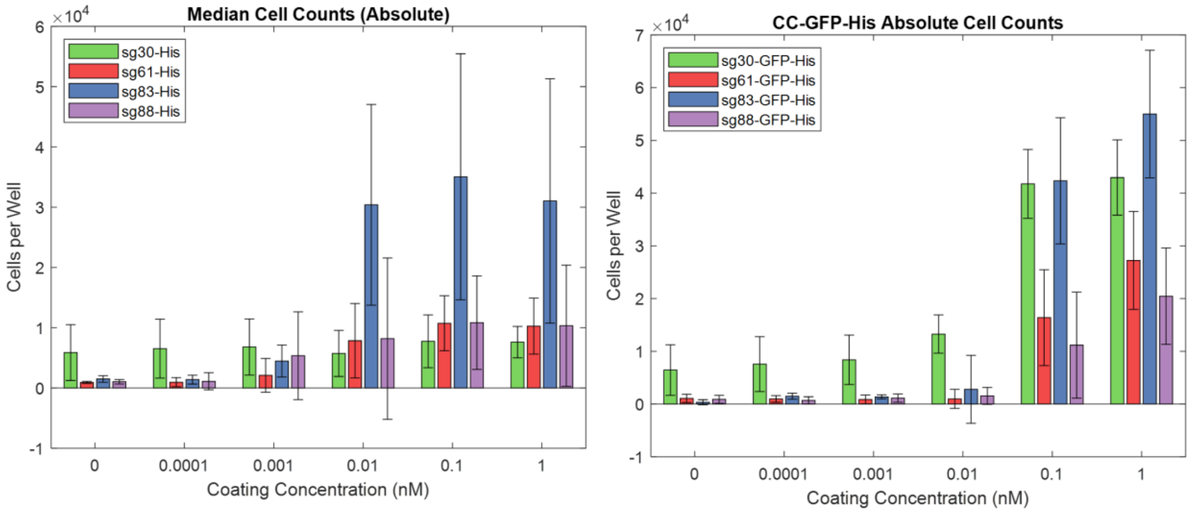


Absolute cell counts for gradient of CC-His and CC-GFP-His cell patterning. Concentrations used for CC-His were 0, 0.01, 0.1, 1, 10, and 100nM, and for CC-GFP-His were 0, 0.0001, 0.001, 0.01, 0.1, and 1nM (5 order of magnitude each). CC-GFP-His concentrations were constrained by expression level. Significantly more cells bound for CC-GFP-His compared with the CC-His conditions for all CCs. Additionally, sg61$^{eGFP}$ cells, corresponding to the sg30-His and sg30-GFP-His conditions, had higher background binding, even for the no CC condition. This is likely due to the His tag on the sg61 construct binding to the $Ni^{2+}$ coating, despite the presence of 20mM imidazole to inhibit this interaction. N=4, error bars are S.D.

S16 – Dot-shaped patterning of K562 helixCAM cells using CC-GFP-His

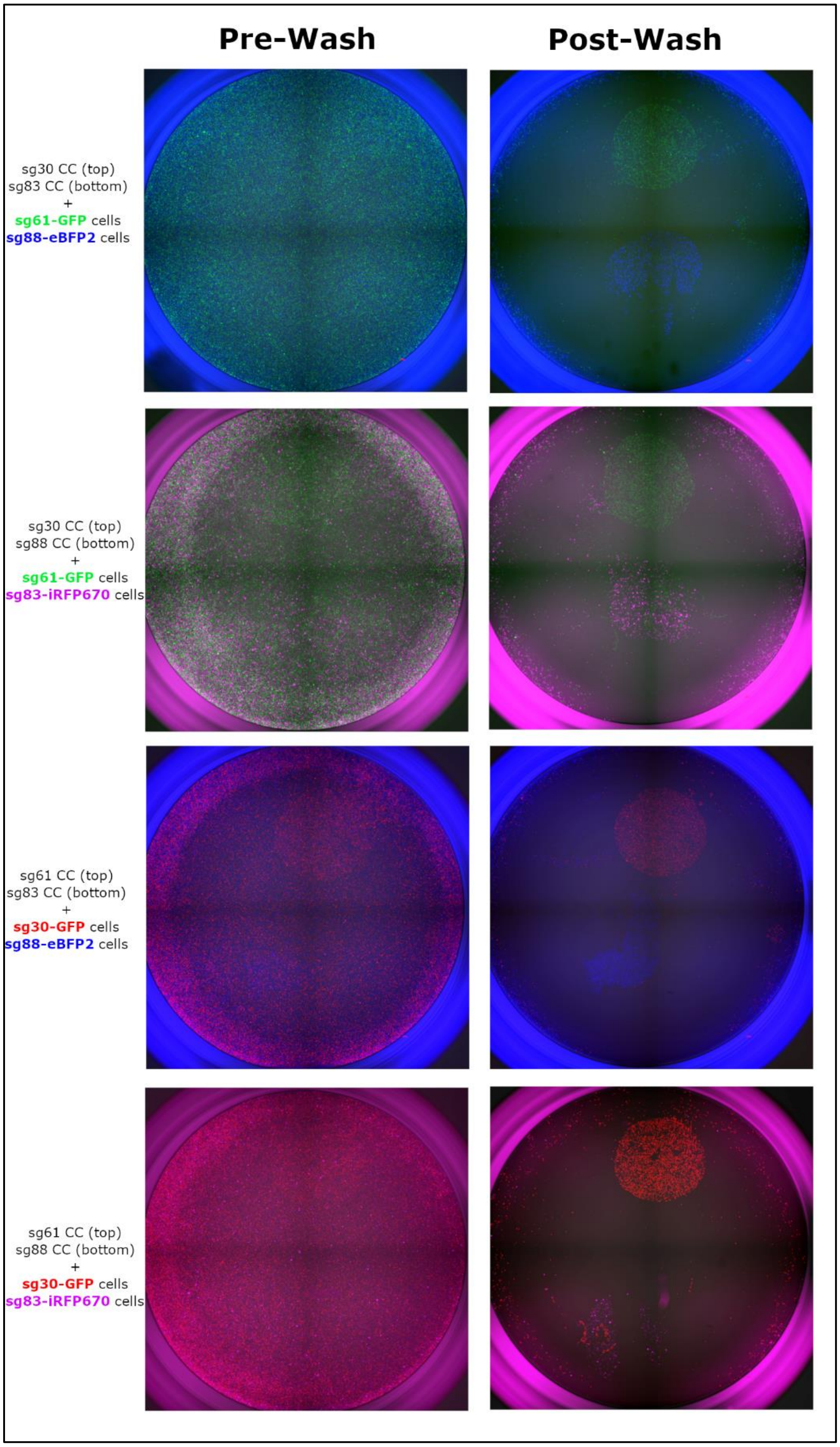

Similar to Figure 7c, Nickel-coated plates were pre-patterned with two distinct Coiled-Coil-GFP-His protein solutions. In this case, 1uL of each protein solution was dotted on either the top of the bottom of the well. The solution was incubated for 30 minutes, washed with PBS+imidazole, then blocked for 30 minutes (also with PBS+imidazole). After removing the blocking buffer, a mixture of the two corresponding K562 helixCAM cells was added into a dual induction/blocking media (1e5 of each cell type in 150uL of DMEM+10%FBS+1%P/S+1ug/mL doxycycline+20mM imidazole). The wells were incubated with the cells for 2 hours, followed by two washes with the same media, and the well was imaged as a 2x2 stitching at 4X magnification. It is possible to see the distinct localization of densely bound cells to the two patterned dots, with few cells in the surrounding region. Since all aspiration and addition of media is done along the bottom of the well, the pipette tips appear to have removed areas of bound cells, but the differential patterning is still clearly visible.

S17 – Table of Key Protein Sequences

| Name | Amino Acid Sequence |
|---|---|
| Z17 domain | NEKEELKSKKAELRNRIEQL<br>KQKREQLKQKIANLRKEIEA<br>YKLQV |
| Z17 mammalian helixCAM | METDTLLLWVLLLWVPGSTG<br>DYPYDVPDYAGANEKEELKS<br>KKAELRNRIEQLKQKREQLK<br>QKIANLRKEIEAYKLQVDEQ<br>KLISEEDLNAVGQDTQEVIV<br>VPHSLPFKVVVISAILALVV<br>LTIISLIILIMLWQKKPR* |
| Z18 domain | SIAATLENDLARLENENARL<br>EKDIANLERDLAKLEREEAY<br>FLQV |
| Z18 mammalian helixCAM | METDTLLLWVLLLWVPGSTG<br>DYPYDVPDYAGASIAATLEN<br>DLARLENENARLEKDIANLE<br>RDLAKLEREEAYFLQVDEQK<br>LISEEDLNAVGQDTQEVIVV<br>PHSLPFKVVVISAILALVVL<br>TIISLIILIMLWQKKPR* |
| P3 domain | SPEDEIQQLEEEIAQLEQKN<br>AALKEKNQALKYG |
| P3 mammalian helixCAM | METDTLLLWVLLLWVPGSTG<br>DYPYDVPDYAGASPEDEIQQ<br>LEEEIAQLEQKNAALKEKNQ<br>ALKYGDEQKLISEEDLNAVG<br>QDTQEVIVVPHSLPFKVVVI<br>SAILALVVLTIISLIILIML<br>WQKKPR* |
| AP4 domain | SPEDELAANEEELQQNEQKL<br>AQIKQKLQAIKYG |
| AP4 mammalian helixCAM | METDTLLLWVLLLWVPGSTG<br>DYPYDVPDYAGASPEDELAA<br>NEEELQQNEQKLAQIKQKLQ<br>AIKYGDEQKLISEEDLNAVG<br>QDTQEVIVVPHSLPFKVVVI<br>SAILALVVLTIISLIILIML<br>WQKKPR* |
| P9 domain | SPEDENQALEQKNAQLKQEI<br>AALEQEIAQLEYG |
| P9 mammalian helixCAM | METDTLLLWVLLLWVPGSTG<br>DYPYDVPDYAGASPEDENQA<br>LEQKNAQLKQEIAALEQEIA |

| | |
|---|---|
| | QLEYGDEQKLISEEDLNAVG<br>QDTQEVIVVPHSLPFKVVVI<br>SAILALVVLTIISLIILIML<br>WQKKPR* |
| AP10 domain | SPEDKLAQIKEKLQQIKEEL<br>AANEEKLQANKYG |
| AP10 mammalian helixCAM | METDTLLLWVLLLWVPGSTG<br>DYPYDVPDYAGASPEDKLAQ<br>IKEKLQQIKEELAANEEKLQ<br>ANKYGDEQKLISEEDLNAVG<br>QDTQEVIVVPHSLPFKVVVI<br>SAILALVVLTIISLIILIML<br>WQKKPR* |
| sGFP30 domain | SPEDEIQALEQENAQLEQKI<br>AALKQENAQLEQEIQALEQ |
| sGFP30 mammalian helixCAM | METDTLLLWVLLLWVPGSTG<br>DGSHHHHHHGASPEDEIQAL<br>EQENAQLEQKIAALKQENAQ<br>LEQEIQALEQDEQKLISEED<br>LNAVGQDTQEVIVVPHSLPF<br>KVVVISAILALVVLTIISLI<br>ILIMLWQKKPR* |
| sGFP61 domain | SPEDKIQALKQKNAQLKQEN<br>AALEQKNAQLKQEIQALEQ |
| sGFP61 mammalian helixCAM | METDTLLLWVLLLWVPGSTG<br>DGSHHHHHHGASPEDKIQAL<br>KQKNAQLKQENAALEQKNAQ<br>LKQEIQALEQDEQKLISEED<br>LNAVGQDTQEVIVVPHSLPF<br>KVVVISAILALVVLTIISLI<br>ILIMLWQKKPR* |
| sGFP83 domain | SPEDKNEELKSKIAELKNKN<br>EALEQKIEQLKQEIANLEK |
| sGFP83 mammalian helixCAM | METDTLLLWVLLLWVPGSTG<br>DGSHHHHHHGASPEDKNEEL<br>KSKIAELKNKNEALEQKIEQ<br>LKQEIANLEKDEQKLISEED<br>LNAVGQDTQEVIVVPHSLPF<br>KVVVISAILALVVLTIISLI<br>ILIMLWQKKPR* |
| sGFP88 domain | SPEDKLATIKNKLAEIKNEL<br>ARNKKELANIERELAKNER |
| sGFP88 mammalian helixCAM | METDTLLLWVLLLWVPGSTG<br>DGSHHHHHHGASPEDKLATI<br>KNKLAEIKNELARNKKELAN<br>IERELAKNERDEQKLISEED |

| | |
|---|---|
| | LNAVGQDTQEVIVVPHSLPF<br>KVVVISAILALVVLTIISLI<br>ILIMLWQKKPR* |
| Sg30-CC | MASPEDEIQALEQENAQLEQ<br>KIAALKQENAQLEQEIQALE<br>QGGGSGGGSGGGSHHHHHH |
| Sg30-GFP-CC | MSPEDEIQALEQENAQLEQK<br>IAALKQENAQLEQEIQALEQ<br>GGSGGGENLYFQSGGSGGSV<br>SKGEELFTGVVPILVELDGD<br>VNGHKFSVSGEGEGDATYGK<br>LTLKFICTTGKLPVPWPTLV<br>TTLTYGVQCFSRYPDHMKQH<br>DFFKSAMPEGYVQERTIFFK<br>DDGNYKTRAEVKFEGDTLVN<br>RIELKGIDFKEDGNILGHKL<br>EYNYNSHNVYIMADKQKNGI<br>KVNFKIRHNIEDGSVQLADH<br>YQQNTPIGDGPVLLPDNHYL<br>STQSALSKDPNEKRDHMVLL<br>EFVTAAGITLGMDELYKHHH<br>HHH |
| Sg61-CC | MASPEDKIQALKQKNAQLKQ<br>ENAALEQKNAQLKQEIQALE<br>QGGGSGGGSGGGSHHHHHH |
| Sg61-GFP-CC | MSPEDKIQALKQKNAQLKQE<br>NAALEQKNAQLKQEIQALEQ<br>GGSGGGENLYFQSGGSGGSV<br>SKGEELFTGVVPILVELDGD<br>VNGHKFSVSGEGEGDATYGK<br>LTLKFICTTGKLPVPWPTLV<br>TTLTYGVQCFSRYPDHMKQH<br>DFFKSAMPEGYVQERTIFFK<br>DDGNYKTRAEVKFEGDTLVN<br>RIELKGIDFKEDGNILGHKL<br>EYNYNSHNVYIMADKQKNGI<br>KVNFKIRHNIEDGSVQLADH<br>YQQNTPIGDGPVLLPDNHYL<br>STQSALSKDPNEKRDHMVLL<br>EFVTAAGITLGMDELYKHHH<br>HHH |
| Sg83-CC | MASPEDKNEELKSKIAELKN<br>KNEALEQKIEQLKQEIANLE<br>KGGGSGGGSGGGSHHHHHH |
| Sg83-GFP-CC | MSPEDKNEELKSKIAELKNK<br>NEALEQKIEQLKQEIANLEK |

| | |
|---|---|
| | GGSGGGENLYFQSGGSGGSV<br>SKGEELFTGVVPILVELDGD<br>VNGHKFSVSGEGEGDATYGK<br>LTLKFICTTGKLPVPWPTLV<br>TTLTYGVQCFSRYPDHMKQH<br>DFFKSAMPEGYVQERTIFFK<br>DDGNYKTRAEVKFEGDTLVN<br>RIELKGIDFKEDGNILGHKL<br>EYNYNSHNVYIMADKQKNGI<br>KVNFKIRHNIEDGSVQLADH<br>YQQNTPIGDGPVLLPDNHYL<br>STQSALSKDPNEKRDHMVLL<br>EFVTAAGITLGMDELYKHHH<br>HHH |
| Sg88-CC | MASPEDKLATIKNKLAEIKN<br>ELARNKKELANIERELAKNE<br>RGGGSGGGSGGGSHHHHHH |
| Sg88-GFP-CC | MSPEDKLATIKNKLAEIKNE<br>LARNKKELANIERELAKNER<br>GGSGGGENLYFQSGGSGGSV<br>SKGEELFTGVVPILVELDGD<br>VNGHKFSVSGEGEGDATYGK<br>LTLKFICTTGKLPVPWPTLV<br>TTLTYGVQCFSRYPDHMKQH<br>DFFKSAMPEGYVQERTIFFK<br>DDGNYKTRAEVKFEGDTLVN<br>RIELKGIDFKEDGNILGHKL<br>EYNYNSHNVYIMADKQKNGI<br>KVNFKIRHNIEDGSVQLADH<br>YQQNTPIGDGPVLLPDNHYL<br>STQSALSKDPNEKRDHMVLL<br>EFVTAAGITLGMDELYKHHH<br>HHH |
